# Convergent and distributed effects of the schizophrenia-associated 3q29 deletion on the human neural transcriptome

**DOI:** 10.1101/2020.05.25.111351

**Authors:** Esra Sefik, Ryan H. Purcell, The Emory 3q29 Project, Elaine F. Walker, Gary J. Bassell, Jennifer G. Mulle

**Author notes:** These authors contributed equally (Esra Sefik, Ryan H. Purcell). Correspondence: Jennifer G. Mulle, MHS, PhD, Emory University School of Medicine, 615 Michael St., Suite 301, Atlanta, Georgia 30322, /.

## Abstract

The 3q29 deletion (3q29Del) confers >40-fold increased risk for schizophrenia. However, no single gene in this interval is definitively associated with disease, prompting the hypothesis that neuropsychiatric sequelae emerge upon loss of multiple functionally-connected genes. 3q29 genes are unevenly annotated and the impact of 3q29Del on the human neural transcriptome is unknown. To systematically formulate unbiased hypotheses about molecular mechanisms linking 3q29Del to neuropsychiatric illness, we conducted a systems-level network analysis of the non-pathological adult human cortical transcriptome and generated evidence-based predictions that relate 3q29 genes to novel functions and disease associations. The 21 protein-coding genes located in the interval segregated into seven clusters of highly co-expressed genes, demonstrating both convergent and distributed effects of 3q29Del across the interrogated transcriptomic landscape. Pathway analysis of these clusters indicated involvement in nervous-system functions, including synaptic signaling and organization, as well as core cellular functions, including transcriptional regulation, post-translational modifications, chromatin remodeling and mitochondrial metabolism. Top network-neighbors of 3q29 genes showed significant overlap with known schizophrenia, autism and intellectual disability-risk genes, suggesting that 3q29Del biology is relevant to idiopathic disease. Leveraging “guilt by association”, we propose nine 3q29 genes, including one hub gene, as prioritized drivers of neuropsychiatric risk. These results provide testable hypotheses for experimental analysis on causal drivers and mechanisms of the largest known genetic risk factor for schizophrenia and highlight the study of normal function in non-pathological post-mortem tissue to further our understanding of psychiatric genetics, especially for rare syndromes like 3q29Del, where access to neural tissue from carriers is unavailable or limited.

## Introduction

Copy number variants (CNVs) offer tractable entry points to investigate the genetic contributions to complex neuropsychiatric diseases. The recurrent 1.6Mb deletion of the 3q29 interval (3q29Del) is robustly associated with schizophrenia spectrum disorders (SZ) [1–4] and is the strongest known risk allele for the disease with an estimated odds ratio >40 [5]. Autism spectrum disorders (ASD) and intellectual disability / developmental delay (IDD) are also enriched in 3q29Del carriers [6–8]. However, it is not currently known which genes within the interval are responsible for the increased neuropsychiatric risk. No single 3q29 interval gene has been definitively associated with SZ, ASD, or ID, prompting the hypothesis that haploinsufficiency of more than one gene is required [9]. The paucity of information regarding the functional roles of most 3q29 interval genes hampers the development of evidence-based hypotheses for deciphering this link. No transcriptomic investigation of 3q29Del in humans has been reported, and it is unclear what impact hemizygous loss of these genes might have in the nervous system.

Among the 21 protein-coding genes of the 3q29Del locus, several have been proposed as drivers of the behavioral phenotypes [10], yet the evidence for their individual association with neuropsychiatric disease remains suggestive. *DLG1* produces a synaptic scaffold protein that interacts with AMPA and NMDA-type glutamate receptors [11–14], the latter of which is hypothesized to be involved in SZ pathogenesis [15]. A *DLG1* polymorphism has been genetically linked to SZ [16, 17]. However, the mouse-specific phenotypes of 3q29Del are not recapitulated by haploinsufficiency of *Dlg1* alone [18]. Another prominent candidate, *PAK2* encodes a brain-expressed protein kinase involved in cytoskeletal dynamics [19] and dendritic spine morphology [20]. Both *DLG1* and *PAK2* are homologous to genes linked to ID [21, 22] and evidence from *Drosophila* indicates that joint haploinsufficiency of both genes simultaneously may be required for synaptic defects rather than either gene individually [23]. A recent study generated select combinatorial knockdowns of 3q29 gene homologs in *Drosophila* and *Xenopus laevis* and proposed that a component of the nuclear cap-binding complex, *NCBP2* [24], mediates neurodevelopmental defects in 3q29Del syndrome [9]. However, in the 3q29Del mouse model, NCBP2 is not decreased at the protein level in brain tissue [18], dampening enthusiasm for this gene as a causal element. It remains unclear which genes and their potential interactions are responsible for 3q29-associated phenotypes.

To avoid annotation-bias [25] and address the knowledge gap for 3q29 genes, we employed a gene co-expression network analysis approach that is rooted in systems biology [26–29], and generated evidence-based predictions that relate individual 3q29 genes to novel functions and disease phenotypes. Accumulating evidence indicates that genes work in conjunction, rather than in isolation, to realize most cellular functions [9, 23, 30]. Genes participating in the same molecular and biological pathways tend to show positively correlated expression with each other (co-expression), as they are often expressed under the control of a coordinated transcriptional regulatory system [31, 32]. In this holistic context, well-characterized genes can be leveraged to infer the functions of understudied genes by studying network patterns that emerge by means of co-expression [33–35]. This *in silico* approach to investigating unknown biology extends the “guilt by association” paradigm [36] that is extensively used for inductive reasoning in other domains to gene-gene interactions in complex biological systems [37–39]. Weighted gene co-expression network analysis (WGCNA) [27, 40] has been successfully deployed to study how genes embedded in network structures jointly affect complex human diseases [41–51]. We employed this paradigm to glean new biological insights into the 3q29Del syndrome.

## Methods and Materials

### The reference dataset

To uncover the network-level operations of genes located in the recurrent 3q29Del locus, we employed WGCNA [27, 40] and organized the non-pathological adult human cortical transcriptome into modules of highly co-expressed genes (Fig. 1a). Given the strong genetic link between 3q29Del and risk for SZ [5], we focused the present network analysis on revealing the clustering patterns of 3q29 interval genes as a function of their expression similarity during adulthood: a period when SZ typically manifests diagnostically, with peak onset in late adolescence and early adulthood [52], and a substantial proportion of patients becoming ill during middle adulthood [53]. A prior study has shown that only 0.7% of genes detected in the neocortex show a temporally regulated profile of differential expression during adulthood (between ages 20-60 years) [54]; hence, gene expression data were pooled across adulthood to derive the present dataset. We further focused our analysis spatially on gene-gene relationships in the PFC: a brain region that subserves a diverse range of cognitive and emotional operations, is most consistently implicated in the etiology of SZ [55] and may be particularly vulnerable to the effects of genetic disruption due to its protracted development [56]. For these reasons, the network was constructed on publicly available transcriptomic data from the Genotype Tissue Expression Project (GTEx) [57], using prefrontal cortex (PFC; Brodmann Area 9) samples from male and female adults (age range = 20-79, 68.2% male) with no known history of psychiatric or neurological disorder (Fig. S1, Table S1.1). Transcriptome profiling was performed by RNA-Seq as described in [57].

**Figure 1.**
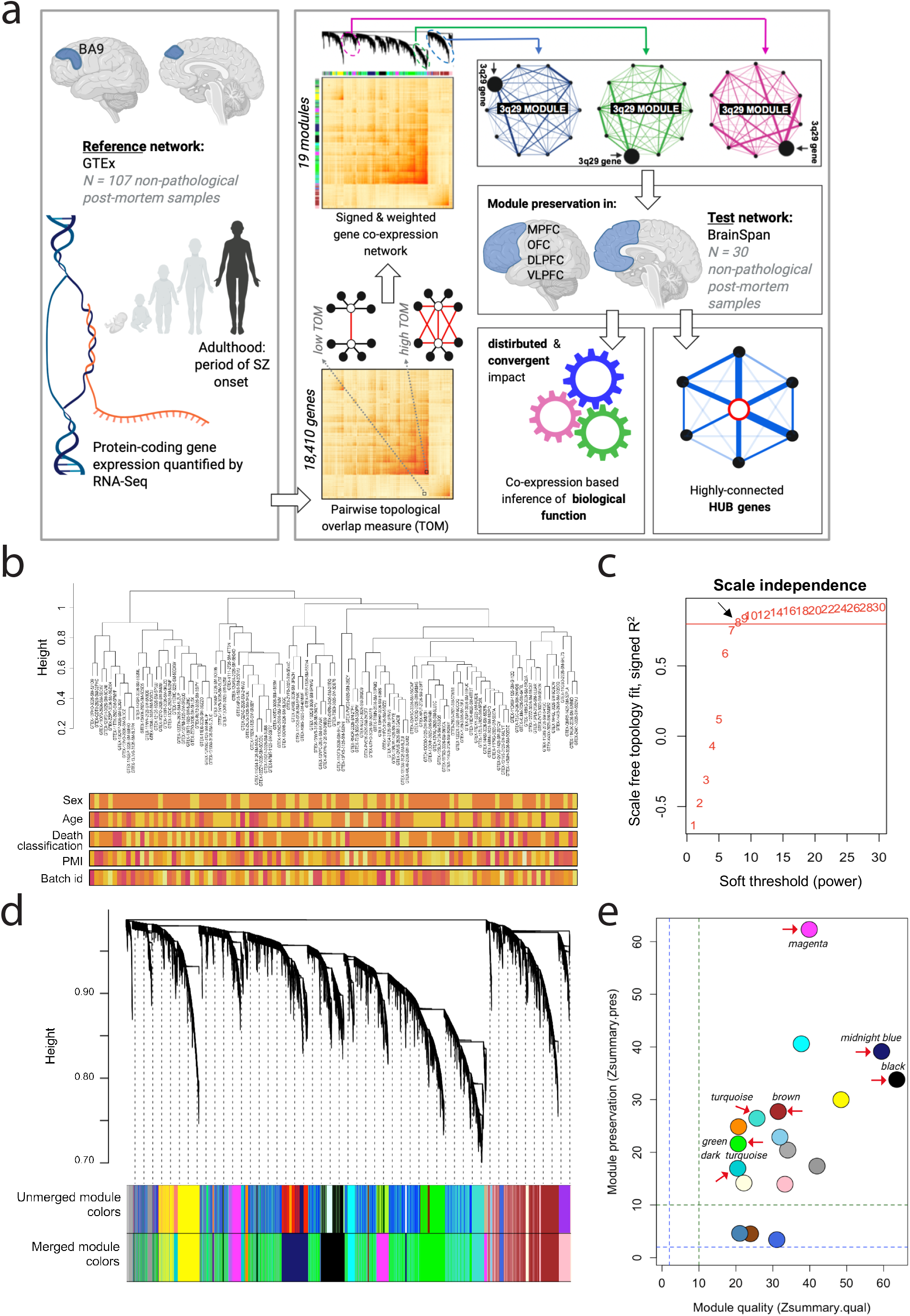
Unbiased weighted gene co-expression network analysis (WGCNA) of the human transcriptome in the healthy adult prefrontal cortex (PFC). **(a)** A schematic of the data analysis workflow underlying WGCNA-derived predictions for functional interrogation of the 3q29Del interval. The reference dataset was obtained from the GTEx Project to construct a systems-level network representation of coordinated gene expression patterns across 107 non-pathological post-mortem samples collected from the Brodmann Area 9 (BA9) of male and female adults with no known history of psychiatric or neurological disorder. Modules of highly co-expressed genes were identified based on their topological overlap measure (TOM). The TOM between two genes is high if the genes have many overlapping network connections, yielding an interconnectedness measure that is proportional to the number of shared neighbors between pairs of genes. The resulting network was screened for modules harboring 3q29 interval genes (3q29 modules), which were then interrogated for biological function and hub genes. A test dataset obtained from the BrainSpan Project was used to validate the reproducibility of this network in an independent sample of 30 non-pathological post-mortem specimens collected from four sub-regions of the PFC from adult males and females with no known history of psychiatric or neurological disorder. These sub-regions are the OFC: orbital frontal cortex, DLPFC: dorsolateral PFC, VLPFC: ventrolateral PFC, and MPFC: medial PFC. **(b)** Sample-level dendrogram and trait heatmaps of the reference dataset. The dendrogram was yielded by hierarchical clustering of 107 GTEx samples using normalized, outlier-removed and residualized gene expression values for 18,410 protein-coding genes. Color bars represent trait heatmaps for sex, age-group (range = 20-79 years), death-classification based on the Hardy scale (range = 0-4), post-mortem interval (PMI) and batch id. The color intensity (from light yellow to red) is proportional to continuous or categorical values (in increasing order) of each variable. For sex, yellow and orange indicate female and male, respectively. Transcriptomic data were corrected for covariance mediated by these variables prior to network construction. Adjusted data reveal no distribution bias associated with the interrogated confounds in sample-level clustering patterns. **(c)** Determination of the soft-thresholding power (*β*) used for WGCNA. A *β* of 8 (black arrow) was identified as the lowest possible power yielding a degree distribution that results in approximate scale-free network topology (SFT R^2^ fit index = 0.8; red line). **(d)** Clustering dendrogram and module assignments of genes, with dissimilarity based on TOM. 18,410 protein coding genes (leaves = genes) clustered into 19 final modules (bottom color bar), detected by the dynamic hybrid tree cut method. Modules with strongly correlated eigengenes (Pearson’s r > 0.8, *P* < 0.05) were amalgamated to eliminate spurious assignment of highly co-expressed genes into separate modules. Color bars reflect module assignments before and after the merging of close modules. **(e)** Composite *Zsummary* scores for module-preservation (how well-defined modules are in an independent test dataset) and module-quality (how well-defined modules are in repeated random splits of the reference dataset). Permutation tests were performed to adjust the observed preservation and quality statistics of each module for random chance by defining Z statistics. All modules (labeled by color) identified in the reference network were preserved (reproducible) in the test network (*Zsummary* > 2; blue line). 15/18 modules, including all 3q29 modules (red arrows), exhibited strong preservation (*Zsummary* > 10; green line). 3/18 modules exhibited moderate preservation (2 < *Zsummary* < 10). All modules demonstrated strong evidence for high quality (*Zsummary* > 10), confirming that the modules identified in the reference network were well-defined and non-random.

Protein-coding transcripts were extracted from the dataset, normalized expression values were log_2_ transformed and summarized at the gene-level, and outlier samples were removed (Fig. S2). The data were corrected for covariance mediated by age, sex, death classification, post-mortem interval (PMI) and batch effect [58, 59] (Fig. 1b, Fig. S1). Genes with zero variance and genes and samples with greater than 50% missing entries (default) were removed [27, 40]. The normalized, outlier-removed, residualized expression values of 18,410 protein-coding genes from 107 samples constituted the final dataset.

### Weighted and signed gene co-expression network construction

The single-block pipeline implemented in the WGCNA R package was employed for network construction [27, 40]. Co-expression similarity was defined by biweight midcorrelation [60, 61]. To capture the continuous nature of interactions and accentuate strong positive correlations, co-expression similarity was transformed into a signed and weighted adjacency matrix by a soft-thresholding procedure that yielded approximate scale-free topology [62–65] (Fig. 1c, Fig. S3). Topological overlap measures (TOM) were calculated from the resulting adjacency matrix to capture not only the univariate correlational relationship between gene-pairs but also the large-scale connections among “neighborhoods” of genes [66, 67] (Fig. 1a). Hence, we measure the interconnectedness of gene-pairs in relation to the commonality of the nodes they connect to.

The modular structure of the data was revealed by average linkage hierarchical clustering on TOM following its transformation into a dissimilarity metric (Fig. 1d). Module definitions used in this study do not use *a priori* knowledge of functionally defined gene-sets. Instead, modules were detected in a data-driven fashion through adaptively pruning the branches of the resulting dendrogram by the dynamic-hybrid-tree-cut method [68]. The expression profile of each identified module was subsequently summarized by a module eigengene (ME) [69], defined as its first principal component. Calculation of MEs amounted to a data reduction method used for effectively correlating entire modules to one another and for establishing the eigengene-based module connectivity measure (kME) of individual genes [40]. To eliminate spurious assignment of genes into separate modules, modules with strongly correlated MEs were amalgamated (Pearson’s r > 0.8, *P* < 0.05, cut height = 0.2) (Fig. 1d).

### Module preservation and quality analyses

To validate the reproducibility of the network modules derived from the GTEx dataset (considered the reference dataset/network), we evaluated module preservation in an independently-ascertained, demographically-comparable transcriptomic dataset, referred to as the test dataset/network (Fig. S4a). For this purpose, publicly available transcriptomic data was obtained from the BrainSpan Developmental Transcriptome Project [54]. 30 non-pathological post-mortem samples from the PFC of male and female adults (age range = 18-37, 50% male) with no known neurological or psychiatric disorder comprised the test dataset (Fig. S4b, Table S1.1). Transcriptome profiling was performed by RNA-Seq as described in [70].

The pre-processed test dataset consisted of normalized and residualized expression values for 18,339 protein-coding genes from 30 samples that were pooled from four sub-regions of the PFC to test whether the co-expression patterns derived from Brodmann Area 9 of the PFC in the reference dataset could be commonly found and robustly defined in broader sampling of tissue across the PFC. Prior to preservation analysis, a sample-level hierarchical clustering of the test dataset was conducted, which revealed no distribution bias associated with PFC sub-region, ruling out tissue of origin as a potential confound in test network construction. (Fig. S4c).

To determine whether properties of within-module topology were preserved in the test network, we calculated a composite, network-based preservation statistic for each module (Z_summary.pres_) by using the *modulePreservation* function of the WGCNA package in R [71]. Z_summary.pres_ is a summary statistic that encompasses multiple density-based and connectivity-based preservation statistics, which are equally important for establishing the overall preservation of a module. To determine whether the observed preservation statistics were higher than expected by chance, we randomly permuted the module assignments in the test dataset (200 times) and derived a standardized Z_summary.pres_ score for each module. To account for this metric’s dependence on module-size, we reduced large modules by randomly sampling 1000 intra-modular nodes. The resulting scores were evaluated according to established thresholds: Z_summary.pres_ < 2, no evidence for preservation; 2 < Z_summary.pres_ < 10, moderate evidence for preservation; Z_summary.pres_ > 10, strong evidence for preservation.

In addition to testing preservation, we measured the quality of the modules that were defined in the reference network by employing a resampling technique that applied the preservation statistics outlined above to repeated random splits of the reference dataset. We assessed the robustness of the identified modules (i.e., how distinct a module is from the background) by calculating a standardized, composite quality statistic (Z_summary.qual_), as described in [71]. The same Z_summary.pres_ thresholds outlined above were used to evaluate Z_summary.qual._

### Functional characterization of network modules harboring 3q29 genes

Pathway analyses of individual modules found to harbor 3q29 genes were performed by g:Profiler (http://biit.cs.ut.ee/gprofiler), using annotations from the Gene Ontology (GO), Reactome and Kyoto Encyclopedia of Genes and Genomes (KEGG) databases. Enriched terms surpassing the adjusted g:SCS significance threshold of *P* < 0.05 were filtered for size and semantic similarity [72].

### Identification of prioritized driver genes and biological mechanisms

Disease-associated genes are often more closely connected to each other than random gene-pairs in a biological network; this non-random network characteristic has enabled the identification of novel genetic risk loci for many diseases [73–77]. To generate data-driven hypotheses about which 3q29 genes are causally linked to the major neuropsychiatric phenotypes of 3q29Del, we tested the overlap between “top neighbors” of individual 3q29 genes and known risk genes for SZ and related disorders (Fig. 3a). A top neighbor was defined as any node whose gene expression profile has a moderate-to-high correlation (Spearman’s rho (*ρ*) ≥ 0.5), *P* < 0.05) with a given 3q29 gene (considered a “seed”) within the same module. Hence, top neighbors were identified by a hard-thresholding method applied only to intramodular edges of a seed that were initially defined by the topological overlap principle.

Lastly, to formulate testable hypotheses about key biological mechanisms that link the 3q29 locus to neuropsychiatric disease, we interrogated the functional enrichment of prioritized driver genes, using the same pathway analysis approach applied to modules. See Supplemental Methods for details.

## Results

### Unbiased gene co-expression network analysis reveals convergent and distributed effects of 3q29 interval genes across the adult human cortical transcriptome

Applying an unsupervised WGCNA approach [27, 40] to publicly-available data from the GTEx Project [57] revealed that the protein-coding transcriptome of the healthy adult human PFC can be organized into a gene co-expression network of 19 modules (labeled by color) (Fig. 1d, Fig. 2a). The identified modules group genes with highly similar expression profiles and likely represent shared function and/or co-regulation. The resulting module sizes ranged from 43 (steel-blue module) to 4,746 (green module) genes, with an average module size of 1,014 genes. To obtain high-quality module definitions, one module (grey module) was reserved for genes that could not be unequivocally assigned to any module. Thus, the grey module was excluded from downstream analysis. The resulting set of modules was screened for membership of 3q29 genes; modules that were found to harbor at least one 3q29 gene are rereferred to as “3q29 modules”.

**Figure 2.**
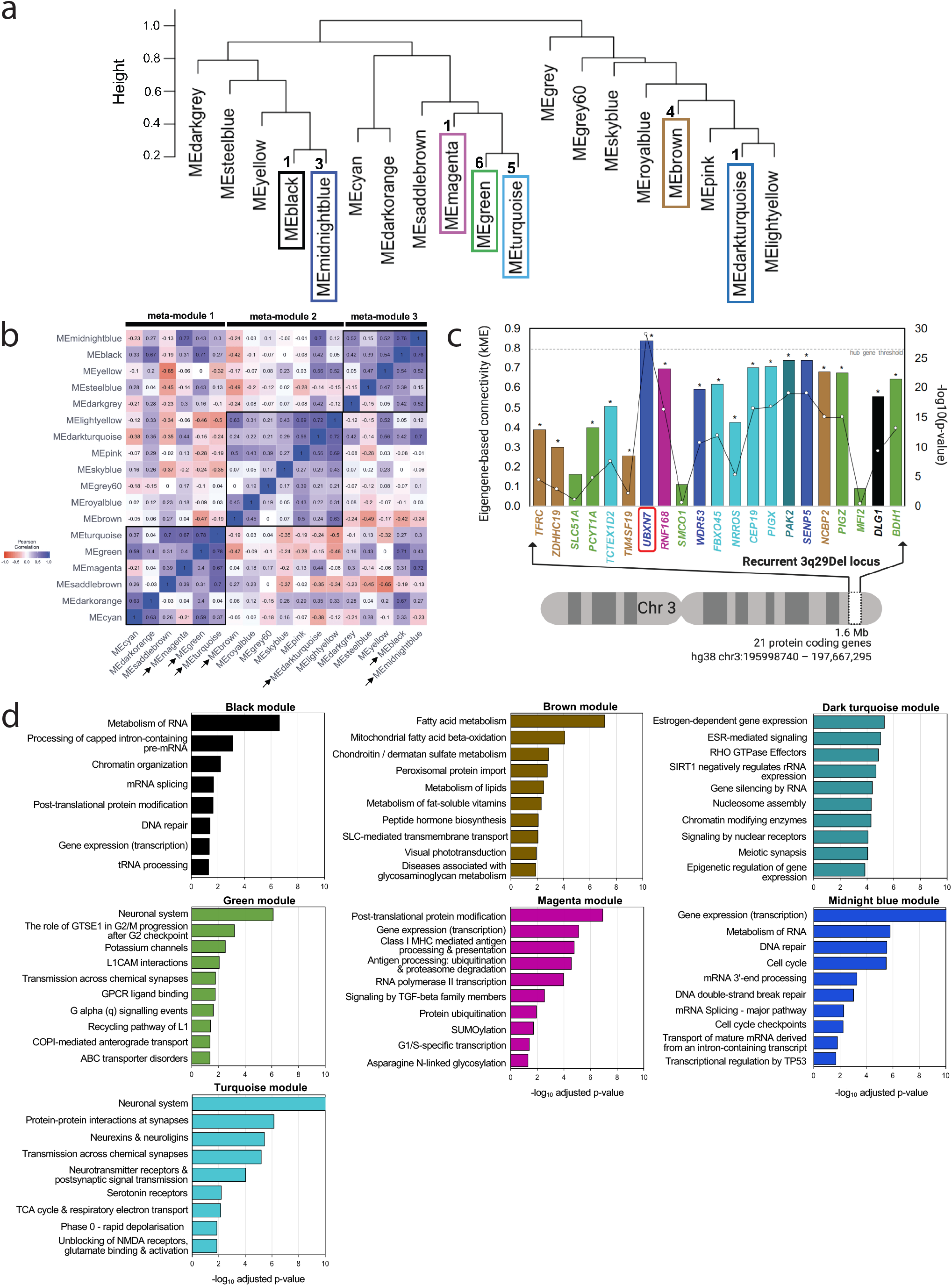
Network-based inference of the functional impact of 3q29Del on the adult human prefrontal cortex (PFC). **(a)** Hierarchical clustering of module eigengenes (ME) that summarize the 19 modules identified by WGCNA. The 21 protein-coding genes located in the recurrent 3q29Del locus were found to be distributed into 7 co-expression modules (3q29 modules; framed), The numbers next to dendrogram branches indicate the total number of 3q29 interval genes found in each 3q29 module. **(b)** Heatmap representing the strength of Pearson’s correlation (*r*) between ME-pairs. The seven 3q29 modules (arrows) further clustered into three higher-level meta-modules, corresponding to squares of blue color (high positive correlation) along the diagonal, also detected as major dendrogram branches in (a). **(c)** Eigengene-based connectivity strength (kME; y-axis) of 3q29 interval genes (x-axis; in chromosomal order) within their respective modules. kME is defined as the Pearson’s correlation between a query gene and a given ME. The line graph indicates the −log10(*P-value*) of the plotted correlation coefficients (z-axis); the asterisks above the graph indicate *P* < 0.05. kME > 0.8 (*P* < 0.05; dotted line) indicates hub (highly connected) gene status. *UBXN7* (red frame) was found to be the only hub gene (kME > 0.8, *P* = 8.33E-30) within the 3q29Del locus. *SMCO1 (*kME = 0.11, *P* = 0.25*), SLC51A (*kME = 0.17, *P* = 0.09) and *MFI2* (kME = 0.09, *P* = 0.35) were found to have non-significant kMEs (P > 0.05) for their respective modules, suggesting peripheral membership. Color indicates module label. **(d)** Top 10 biological pathways (Reactome database) significantly enriched in 3q29 modules (adjusted-*P* < 0.05; capped at −log10 (adjusted-*P* = 10)). The g:SCS method was used for multiple testing correction. The observed enrichment profile of the queried modules for known biological processes and pathways indicates that genes co-clustering in 3q29 modules show coordinated expression and converge upon overlapping biological functions, more than expected by chance. The functional associations of gene-sets comprising individual 3q29 modules were leveraged to infer likely molecular consequences of 3q29Del in the adult human PFC.

To ensure the reproducibility and robustness of our network analysis results, we tested the preservation and quality of the identified modules in an independent dataset obtained from the BrainSpan Developmental Transcriptome Project [54] (Fig. S4) and in repeated random splits of the reference dataset. These analyses revealed the replicable, well-defined and non-random nature of the identified network modules (Fig. 1d, Fig. S5, Table S1.4).

The 21 protein-coding genes located in the 3q29 interval clustered into seven modules (Fig. 2a): black (one 3q29 gene), brown (four 3q29 genes), dark-turquoise (one 3q29 gene), green (six 3q29 genes), magenta (one 3q29 gene), midnight-blue (three 3q29 genes), and turquoise (five 3q29 genes). Within this network, 18 (86%) of the 3q29 interval genes concentrate into just four modules (Fig. 2a), suggesting that the haploinsufficiency of sets of genes within the locus may perturb the same biological processes via multiple hits, cumulatively disrupting redundancy and compensatory resiliency in the normative regulation of cellular functions.

To evaluate whether modules further clustered within larger meta-modules that represent the higher-order organization of the transcriptome, we identified meta-modules as tight clusters of positively correlated MEs, detectable as major branches of the eigengene dendrogram [78] (Fig. 2a, 2b). Meta-modules were screened to identify the grouping patterns among 3q29 modules, allowing exploration of extra-modular interactions. This analysis revealed that the 3q29 modules further cluster into three higher level meta-modules (Fig. 2b), which likely reflect dependencies and interactions between pathways involving 3q29 genes. Simultaneously, leading presumptive candidates *DLG1* and *PAK2* were found in opposite branches of the network, demonstrating the distributed effects of this CNV across the transcriptomic landscape.

### Pathway analysis points to functional involvement of the 3q29 locus in nervous-system functions and core aspects of cell biology

Since highly co-expressed genes often share similar functions [31, 32], biological processes and pathways that are enriched in a co-expression module can be used to infer functional information for poorly annotated genes of that module. Functional enrichment analysis of 3q29 modules showed that their constituent genes converge onto canonical biological processes at proportions greater than expected by chance, indicating that these modules are biologically relevant units (Fig. 2d, Table S1.5). The turquoise and green modules showed overrepresentation of roles specific to the neuronal system and implicate involvement in multiple synaptic properties. Other 3q29 modules point to biological pathways that may also underlie neuropsychiatric pathology in 3q29Del. The magenta module was predominantly enriched for protein modification, turnover, and localization. Additionally, a link was identified between the magenta module and the initiation of major histocompatibility complex class-I (MHC-I)-dependent immune responses, driven by a genomic locus implicated in the etiology of SZ [79, 80]. On the other hand, overrepresented pathways in the black module encompass regulation of gene expression and maintenance of the integrity of the cellular genome, including DNA repair, and the metabolism and processing of RNA. The midnight-blue module shared enriched roles with the black module, validating their shared meta-module structure, yet it was set apart by its involvement in cell cycle regulation. The brown module revealed primary enrichment for cellular metabolism and mitochondrial function, whereas the dark-turquoise module coalesced genes involved in epigenetic mechanisms and in signal transduction pathways mediated by Rho GTPases. This latter function may be attributable to *PAK2*, which encodes a known Rho GTPase effector. Intriguingly, this module was also enriched for estrogen receptor-mediated signaling, suggesting a potential mechanism for sex-specific effects. Taken together, functional characterization of the 3q29 modules point to novel mechanisms of shared or synchronized action for co-clustering 3q29 genes (Fig. 2d).

### UBXN7 is a highly connected cortical hub-gene predicted to play a crucial role in the neuropsychiatric sequela of 3q29Del

Targeted disruption of a highly-connected “hub” gene produces a more deleterious effect on network function and yields a larger number of phenotypic outcomes than randomly selected or less connected genes [81, 82]. Hence, we sought to measure how strongly connected individual 3q29 genes are to their modules by evaluating their intramodular kME (Fig. 2c, Table S1.3), defined as the Pearson’s correlation between the expression profile of a gene and the eigengene of its assigned module [27, 40, 69].

Genes with high intramodular kME are considered hub genes that are predicted to be critical components of the overall function of their module [27]. Nodes with high intramodular kME often have high intramodular connectivity (kIM), which reflects sum of adjacencies to other nodes [40]. However, an advantage of using kME over other connectivity metrics, is its defined *P*-value and values that lie between −1 and 1, allowing comparison across modules that differ in size. To generate rigorous predictions about which 3q29 genes, if any, are intramodular hub genes, we adopted a conservative criterion that defines hub genes as nodes with kME > 0.8 (P < 0.05). Only one 3q29 interval gene was identified as a hub gene: *UBXN7* (kME = 0.84, *P* = 8.33E-30), which encodes a ubiquitin ligase-substrate adaptor [83, 84].

*SMCO1, SLC51A*, and *MFI2* had non-significant kMEs (*P* < 0.05) for their module, suggesting low intramodular connectivity. These 3q29 genes are detected but display very low abundance in the human cerebral cortex [85] (Table S1.6), which may relate to their peripheral network assignments in our analysis. Consequently, *SMCO1, SLC51A* and *MFI2* were excluded from downstream analysis to derive the most parsimonious prioritization of driver genes based on tight network connections. A complete list of gene-sets for each module and kME values are provided in Tables S1.2-3.

### Nine 3q29 interval genes form transcriptomic subnetworks enriched for known SZ, ASD and IDD-risk genes

We next identified a refined subset of target genes (top neighbors) that not only co-cluster based on TOM but also have a strong pairwise correlation with 3q29 genes (Fig. 3a). Several 3q29 genes were found to be top neighbors of one another. *FBXO45* (*ρ* = 0.5, *P* = 5.43E-09) and *PIGX* (*ρ* = 0.6, *P* = 1.24E-10) are top-neighbors of *CEP19*, while *SENP5* and *WDR53* are top-neighbors of each other *(ρ* = 0.5, *P* = 1.05E-07) (Table S2.1). On the other hand, intramodular connections of *TM4SF19* and *ZDHHC19* did not meet top neighborhood criteria; hence they were not included in downstream analysis. Similar to *SMCO1, SLC51A* and *MFI2*, their mRNA expression profiles indicate low abundance in the cerebral cortex (Table S1.6), which likely reflects their lack of strong network connections.

**Figure 3.**
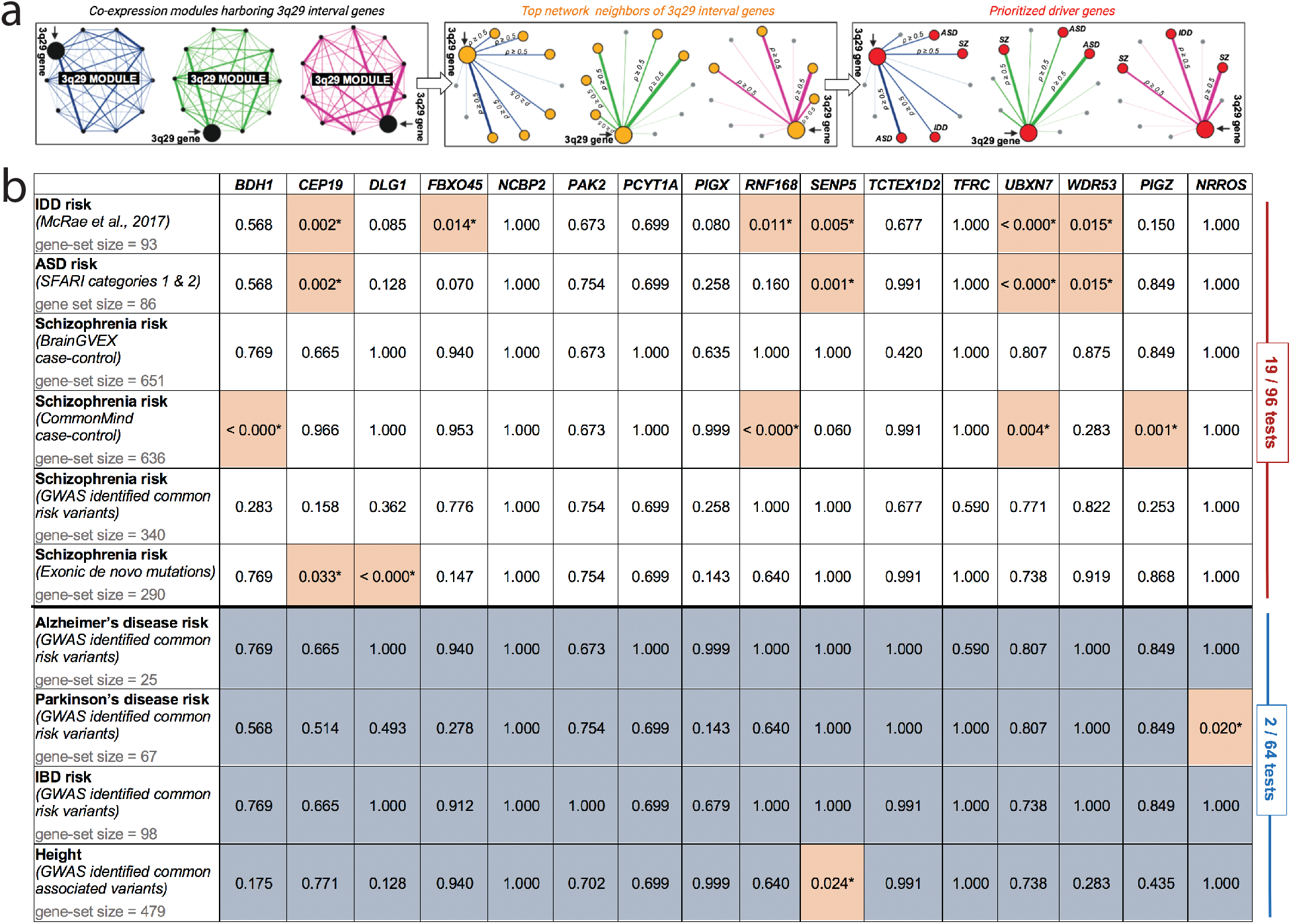
3q29 interval genes form transcriptomic subnetworks enriched for known schizophrenia, autism and intellectual / developmental disability-risk genes. (**a**) Schematic of strategy to test the neuropsychiatric disease burden associated with top network neighbors of 3q29 interval genes and to refine a list of prioritized driver genes. To minimize false positives, 3q29 modules were reduced to strongly connected top neighbors (yellow nodes) of individual 3q29 genes, which were then screened for a significant overlap with known risk genes (red nodes) for schizophrenia (SZ), autism spectrum disorders (ASD), and intellectual / developmental disability (IDD), spanning known associations over a wide spectrum of allele frequencies. A top neighbor was defined as any node whose gene expression profile had a moderate-to-high pairwise correlation (Spearman’s rho (*ρ)* ≥ 0.5, *P* < 0.05) with a 3q29 interval gene within the same module. By leveraging the guilt by association principle, the 3q29 interval genes that showed a significant enrichment of known SZ, ASD, and/or IDD risk genes among their respective top neighbors were prioritized as likely drivers of the neurodevelopmental and psychiatric consequences of 3q29Del, along with their SZ, ASD, and/or IDD-related top neighbors from significant overlap tests. (**b**) Adjusted p-values from hypergeometric tests identifying the significance of the overlap between top neighbors of individual 3q29 genes and known risk genes for SZ, ASD, and IDD. Risk gene-sets for three traits with no known association to the 3q29Del syndrome were also tested for over-representation as negative controls. Common variants associated with height were included as another negative control to rule out a potential bias introduced by gene-set size. Nine protein-coding genes from the 3q29 interval formed transcriptomic subnetworks that are significantly enriched for known SZ, ASD and/or IDD risk genes (orange highlight, adjusted-*P* < 0.05). The proportion of hypergeometric tests with significant overrepresentation of SZ, ASD, and IDD gene-sets (19/96) was found to be an order of magnitude larger than that of the negative control tests (2/64), demonstrating the high specificity of the identified enrichment patterns for reported 3q29Del-associated phenotypes.

The human transcriptome is theorized to demonstrate non-random topological characteristics, where disease genes interact with other disease genes that underlie a common pathophenotype [86]. Concordant with this prediction, within the top neighbors of 3q29 genes, we found several genes that have been extensively implicated in neuropsychiatric disease (Table S2.1). These include *MECP2*, *NRXN1*, *GRIN2A*, *GRIN2B*, *CHD8*, *SATB2*, *CNTNAP2*, *FOXP1*, *PTEN*, and *SCN2A*. Motivated by this observation, we asked whether top neighbors of individual 3q29 genes significantly overlap with known SZ, ASD or intellectual disability / developmental delay (IDD)-risk genes (Fig. 3a). We curated six evidence-based lists of SZ [87–91], ASD [92, 93] and IDD-risk genes [94], which span loci across a wide range of the allele frequency spectrum and include post-mortem findings from case-control gene expression studies (Table S2.2). Hypergeometric tests were conducted to gauge the probability of the overlap between curated gene-sets and top neighbors, as implemented in the *GeneOverlap* package in R [95], followed by Benjamini-Hochberg multiple testing correction. 3q29 genes whose top neighbors showed an overrepresentation of SZ, ASD and/or IDD risk genes (adjusted *P* < 0.05) were subsequently prioritized as likely genetic drivers of neuropsychiatric risk in 3q29Del syndrome, along with their SZ, ASD, and/or IDD-related top neighbors from the enrichment findings (Fig. 3a). We found overrepresentation of one or more established risk gene-sets among the top neighbors of nine 3q29 genes (Benjamini-Hochberg adjusted-*P* < 0.05) (Fig. 3b, Table S2.3).

To evaluate the specificity of the identified patterns of polygenic disease burden, we also tested these top neighbors for overlap with known Parkinson’s disease (PD) [96], late-onset Alzheimer’s disease (AD) [97], and inflammatory bowel disease (IBD) risk genes [98] (Table S2.2). These phenotypes have no known link to 3q29Del, thus, their risk loci were considered negative controls. Common variants associated with height [99] (Table S2.2) were included as a fourth negative control to rule out a potential bias associated with large differences in the sizes of curated gene-sets.

Concurrently, we found no statistically significant evidence for overrepresentation of AD or IBD-risk genes among the interrogated top neighbors (Fig. 3b). Only the top neighbors of *SENP5* showed a significant overlap with height-associated genes (adjusted*-P* = 2.36E-02), and the top neighbors of *NRROS*, which did not show an enrichment for known IDD, ASD, or SZ risk genes, exhibited a small but significant overlap with known PD-risk genes (adjusted*-P* = 2.00E-02) (Fig. 3b). Overall, 19 out of 96 hypergeometric tests (20%) revealed a significant overrepresentation of SZ, ASD, and/or IDD-risk gene-sets among the top neighbors of 3q29 genes. By contrast, only 2 out of 64 (3%) hypergeometric tests indicated a significant overlap with the negative control gene-sets. The substantial margin between these two enrichment ratios supports the high specificity of our network-derived inferences for uncovering biology relevant to 3q29Del. By leveraging guilt by association, we prioritize *BDH1, CEP19, DLG1, FBXO45, PIGZ, RNF168, SENP5, UBXN7, and WDR53*, along with their 284 unique SZ, ASD, and/or IDD-related top neighbors from significant overlap tests as likely drivers of the neuropsychiatric consequences of 3q29Del (Fig. 4a, Table S2.3).

**Figure 4.**
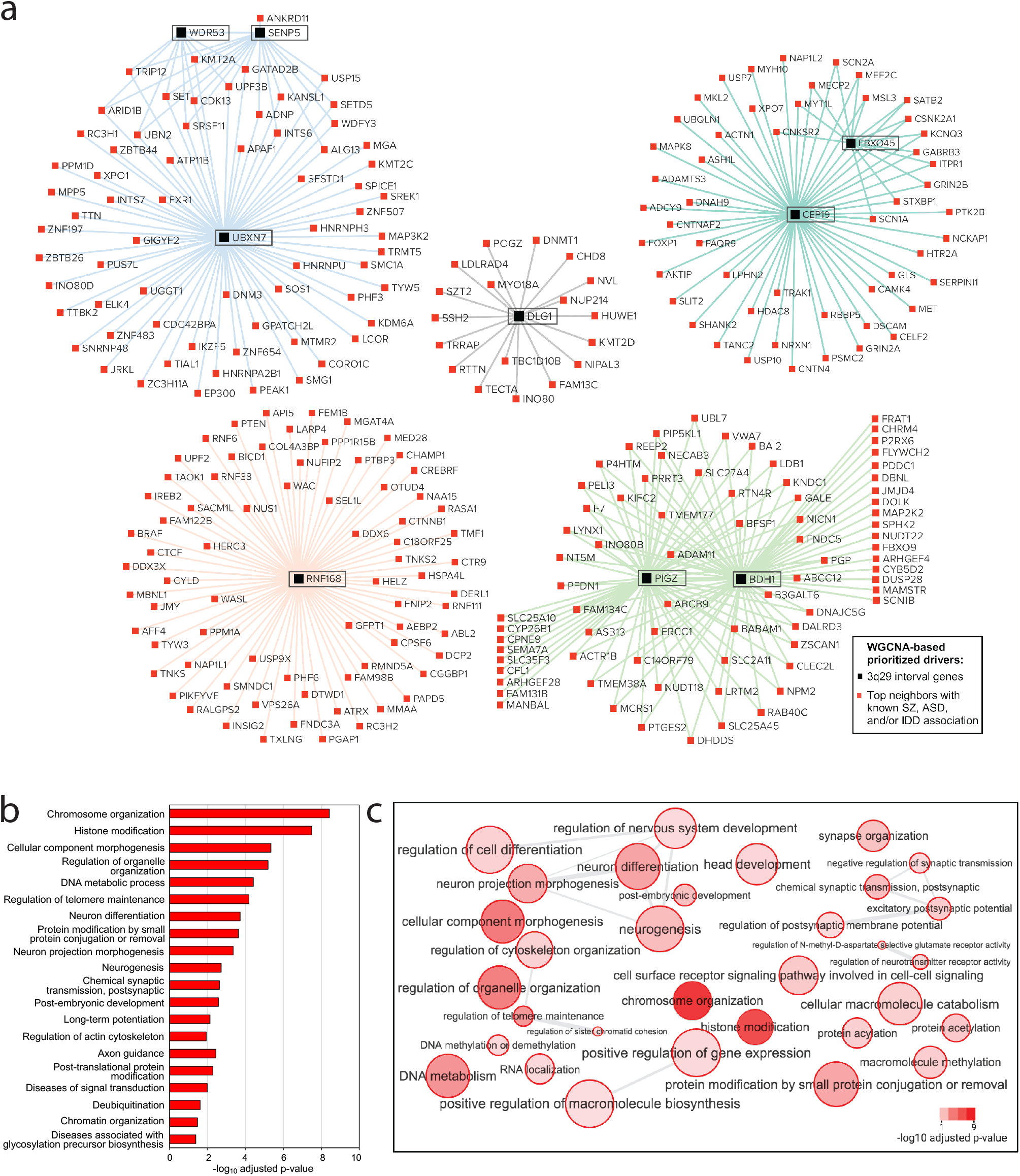
Network of prioritized drivers predicted to contribute to the neuropsychiatric sequelae of 3q29Del. **(a)** Nine protein-coding genes from the 3q29 interval formed top-neighbor-based transcriptomic subnetworks that were significantly enriched for known schizophrenia (SZ), autism spectrum disorder (ASD) and intellectual / developmental disability (IDD)-risk genes (adjusted-*P* < 0.05). Black and red nodes illustrated in this network diagram represent these 9 3q29 genes and their 284 top neighbors with known SZ, ASD and/or IDD-association, respectively. The union of these prioritized genes constitute 293 genetic drivers predicted to contribute to the neurodevelopmental and psychiatric phenotypes of 3q29Del. The color of network edges that connect node-pairs represents module assignment. **(b)** Top 20 biological processes and pathways with significant enrichment among prioritized drivers (adjusted-*P* < 0.05). The identified Gene Ontology biological processes (GO: BP) and Reactome and KEGG biological pathways point to key mechanisms through which select genes within the 3q29Del locus and their likely partners outside the interval are predicted to influence susceptibility to SZ, ASD, and IDD. **(c)** Organization of all statistically significant biological processes enriched in prioritized drivers into a network of related functional annotation categories. GO:BP terms are connected if they have a high overlap (share many genes); edge width represents magnitude of the overlap.

### Disease-relevant driver genes prioritized by network analysis load onto key biological pathways linked to neuropsychiatric disorders

To formulate testable hypotheses about the biological mechanisms linking the 3q29 locus to neuropsychiatric phenotypes, we interrogated whether the prioritized driver genes identified in our network analysis assemble into known biological pathways. Functional enrichment analysis on the union of 293 prioritized driver genes (including nine 3q29 genes) revealed significant overrepresentation of eight biological pathways annotated by the Reactome and KEGG databases (Fig. 4b, Table S2.4). These include axon guidance (adjusted*-P* = 3.64E-03), long-term potentiation (adjusted*-P* = 7.29E-03), and regulation of actin cytoskeleton (adjusted*-P* = 1.17E-02). Additionally, several GO biological processes (GO:BP), including chromosome organization (adjusted*-P* = 3.81E-09), histone modification (adjusted*-P* = 3.31E-08), neuron differentiation (adjusted*-P* = 1.88E-04), neurogenesis (adjusted*-P* = 1.89E-03), and excitatory postsynaptic potential (adjusted*-P* = 8.97E-03) were overrepresented among the predicted drivers (Fig. 4b-c, Table S2.4). We hypothesize that the disruption of one or more of these biological pathways and processes, some of which have been demonstrated to be altered in idiopathic SZ and ASD [2, 100], lie on the casual pathway to neuropsychopathology in 3q29Del syndrome. For extended results, see Supplemental Results.

## Discussion

The 3q29Del has been reliably associated with extraordinary risk for serious neuropsychiatric illness and therefore may offer key insights to advance our understanding of the biological basis of these complex disorders. Currently, the driver genes and affected biological pathways that link 3q29Del to neuropsychiatric pathology remain unknown. To avoid bias introduced by annotation-based criteria in the formulation of mechanistic hypotheses, we engaged a system-level vantage point and interrogated the collective behavior of 3q29 interval genes with the global protein-coding transcriptome of the healthy human brain. We leveraged publicly available transcriptomic data from the GTEx Project [57] to perform WGCNA [27, 40] on postmortem cortical samples from donors with no known history of psychiatric or neurological disease. Given that precise actions of genes are often dependent on tissue and age [54], we focused our analysis on the adult PFC, a brain region most consistently implicated in SZ [55], which typically strikes during adulthood. We analyzed the resulting network to identify the modular properties and undirected connectivity patterns of the 3q29 interval, which yielded key predictions into inter-related functions and disease associations. Our findings provide foundational information to formulate rigorous, targeted and testable hypotheses on the causal drivers and mechanisms underlying the largest known single genetic risk factor for SZ.

Genomic studies have identified several recurrent CNVs that confer high risk for neuropsychiatric disorders [2, 8]. The current challenge is to understand which genes within these loci are the major drivers of risk. In the 3q29 locus, *DLG1* and *PAK2* have been most often proposed as candidate drivers of neuropsychiatric phenotypes [10, 23, 101]. Indeed, a recent literature search revealed more publications related to these genes than of all other 3q29Del genes combined (Fig. S6). Consistent with previous reports of an association between *DLG1* and SZ [16, 17], the current study presents network-level evidence for prioritizing *DLG1* as a neuropsychiatric disease-linked gene. Surprisingly, however, our analysis does not support inclusion of *PAK2* as a predicted driver of neuropsychiatric risk. Instead, our results lend support to *DLG1* and eight other 3q29 genes, most of which are largely understudied, as key players in 3q29Del syndrome. Our unbiased approach prioritizes *BDH1, CEP19*, *DLG1*, *FBXO45, PIGZ, RNF168, SENP5, UBXN7* and *WDR53* as primary drivers.

It is currently unknown whether the biological basis of neuropsychiatric risk associated with recurrent CNVs overlaps with that of individuals who share the same clinical diagnosis but do not share the same rare genetic variant. Our findings suggest that molecular perturbations caused by the hemizygous deletion of select 3q29 genes may overlap with the genetic etiologies contributing to idiopathic forms of SZ, ASD and ID. Disease-relevant driver genes prioritized by our network analysis are enriched for canonical biological pathways, such as neurogenesis, neuron differentiation, synapse organization, excitatory postsynaptic potential, long-term potentiation, axon guidance, regulation of actin cytoskeleton, signal transduction, post-translational protein modifications, chromatin organization and histone modification. We hypothesize that the disruption of one or more of these biological processes, some of which are altered in idiopathic SZ and ASD [2, 100], lie on the casual pathway to neuropsychopathology in 3q29Del syndrome.

No single gene within the interval has been definitively associated with neuropsychiatric disease, prompting the hypothesis that neuropathology in 3q29Del emerges upon loss of multiple genes that are functionally connected. While a single nucleotide polymorphism in *DLG1* has been associated with SZ in a case-control study [16, 17], the risk associated with this variant does not approach that of 3q29Del, suggesting that the neuropsychiatric risk associated with this CNV is distributed across more than one gene in the locus. To investigate functional connections across multiple 3q29 genes, we conducted an unsupervised analysis of the modular organization of the adult human PFC. We found the 21 3q29 genes distributed into three meta-modules and seven modules, with 18 genes converging into just four modules. Hence, 3q29 genes display both distributed and convergent effects in the adult human cortical transcriptome. Rather than functioning as independent agents, sets of 3q29 genes may have shared and/or synchronized function and constitute interacting sources of pathology. It is conceivable that the consequences of the haploinsufficiency arise through the weakening of multiple distinct pathways that normally provide protective redundancies (distributed model), and/or through multiple insults to a functionally connected module that cumulatively disrupt resiliency (convergent biology model). These hypotheses warrant further testing.

A major goal of this study was to infer unknown functions for understudied 3q29 genes by leveraging well-studied co-clustering genes. Pathway analysis of modules harboring 3q29 genes revealed likely functional involvement of the 3q29 locus in not only nervous-system specific functions, but also in core aspects of cell biology that are non-specific to an organ system. The closely-related black and midnight-blue modules were significantly associated with regulation of gene expression, chromatin organization and DNA repair. The green and turquoise modules were both associated with nervous system development and function, and in particular, regulation and organization of synaptic signaling and components. This finding is surprising because most of the 3q29 genes located in these latter two modules have not been identified as synaptic genes. Similarly, the 3q29 genes in the black and midnight-blue modules have not been implicated in gene regulatory pathways or DNA repair. We maintain that biological functions of poorly annotated genes can be inferred through the graph-based modeling of inter-gene relationships. Thereby, we predict novel roles for individual 3q29 genes in functions related to synaptic transmission, modulation of neurotransmission, synapse structure and function, mitochondrial metabolism, transcriptional and translational regulation, chromatin remodeling, cell cycle regulation, and protein modification, localization and turnover. We propose that the subset of predicted functions that are non-specific to an organ system likely contribute to global developmental outcomes in 3q29Del.

Analysis of eigengene-based connectivity revealed that *UBXN7* is a hub gene, with top neighbors enriched for known association with all three major neuropsychiatric phenotypes of 3q29Del. Hub nodes of biological networks are often associated with human disease [102, 103]. Disease-genes, identified from OMIM’s Morbid Map of the Human Genome, disproportionately exhibit hub-gene characteristics, with protein products participating in more known protein-protein interactions than that of non-disease genes [104]. Supported by this literature, we predict that (1) *UBXN7* exerts critical influence on a large network of co-expressed genes, and (2) loss-of-function (LoF) mutations in *UBXN7* can cause major dysfunction in affiliated biological pathways (indeed, its pLI score = 0.99, i.e. extremely intolerant to LoF [105]). We prioritize *UBXN7* as a major driver in 3q29Del syndrome. *UBXN7* has not been previously linked to neuropsychiatric disorders or proposed as a candidate driver of 3q29Del syndrome. However, *UBXN7* has been reported to regulate the ASD-associated E3 ubiquitin ligase Cullin-3 (*CUL3*) [106], an interaction that deserves more attention in light of our findings [84].

One limitation of the current study is its singular focus on protein-coding elements. How the non-coding elements of the interval, along with splice variants, integrate into the predictions formulated in this study is ripe for future investigation. Overall, the transcriptomic network identified in this study is predicted to connect 3q29 interval genes with gene-sets outside the interval that participate in the same or overlapping biological process and associate with similar disease phenotypes. Perturbation of 3q29 interval gene dosage is expected to also perturb the functioning of network-partners outside the recurrent 3q29Del locus. However, note that the underlying structure of weighted gene co-expression networks is agnostic to the mechanistic order of cellular and molecular events. The information necessary to derive the order of biological interactions is not an explicit outcome of gene co-expression itself, since such inferences require time-dependent analysis of combinatorial interactions between nodes. As a result, some of the network partners identified in this study are expected to function upstream of their co-expressed 3q29 interval partner, and likely not be affected by 3q29Del. Moreover, the complex interactions between molecules can be dynamic across time and space [54]. Hence, a future direction will be to ask whether the network connections formed by 3q29 interval genes in the adult PFC show differential expression in the neural tissue of 3q29Del carriers and whether they show temporal and/or spatial variation.

Finally, our analysis does not preclude the possibility that other 3q29 interval genes moderate phenotypic expressivity. For example, while the dark turquoise module (including *PAK2*) did not harbor prioritized driver genes, it was significantly associated with estrogen receptor-mediated signaling. An intriguing emerging feature of 3q29Del syndrome is the markedly reduced sex bias in risk for ASD [7]. Additional studies will be required to assess the drivers of sex-specific phenotypes of 3q29Del syndrome.

Now that recurrent, highly penetrant CNV loci have been identified as important risk factors for neuropsychiatric disorders, determination of the component genes driving this risk is the next step toward deciphering mechanisms. We used an unbiased systems biology approach that leveraged the power of open data to infer unknown functions for understudied 3q29 interval genes, and to refine the 3q29 locus to nine prioritized driver genes, including one hub gene. Importantly, this approach can partially overcome barriers to formulation of relevant hypotheses that are introduced by poor annotation of interval genes, without requiring laborious, expensive, and time-consuming experiments to functionally characterize all genes within the interval. Our results reveal the power of this approach for prioritization of putative drivers. Ongoing and future studies will be directed at understanding how these genes work in concert and how multiple haploinsufficiencies confer risk for neuropsychiatric disease.

## Supporting information

Supplemental Methods and Results

## Acknowledgments

This study was supported by NIH R01 MH110701 (J.G.M. and G.J.B.), F32 MH124273 (R.H.P.), and the Emory University School of Medicine.

The GTEx data (release version 6) used for the analyses described in this manuscript were downloaded from the GTEx portal on 01/08/2019 (http://www.gtexportal.org/home/datasets/, file name: “GTEx_Analysis_v6_RNA-seq_RNA-SeQCv1.1.8_gene_rpkm.gct.gz”); dbGaP accession number: phs000424.v6.p1. The Genotype-Tissue Expression (GTEx) Project was supported by the Common Fund of the Office of the Director of the National Institutes of Health, and by NCI, NHGRI, NHLBI, NIDA, NIMH, and NINDS.

The BrainSpan Developmental Transcriptome dataset used to construct the test network was downloaded from the Allen Brain Atlas portal on 01/12/2019 (https://www.brainspan.org/static/download/, file name: “RNA-Seq Gencode v10 summarized to genes”); dbGaP accession number: phs000755.v2.p1.

The authors gratefully acknowledge the contributions of the members of the Emory 3q29 Project: Katrina Aberizk, Hallie Averbach, T. Lindsey Burrell, Shanthi Cambala, Grace Carlock, Tamara Caspary, Joseph F. Cubells, David Cutler, Paul A. Dawson, Michael P. Epstein, Roberto Espana, Michael J. Gambello, Katrina Goines, Ryan Guest, Henry R. Johnston, Cheryl Klaiman, Sookyong Koh, Elizabeth J. Leslie, Longchuan Li, Bryan Mak, Tamika Malone, Michael Mortillo, Trenell Mosley, Melissa M. Murphy, Derek Novacek, Rebecca M. Pollak, Rossana Sanchez, Celine A. Saulnier, Jason Schroeder, Sarah Shultz, Nikisha Sisodiya, Steven Sloan, Stephen T. Warren, David Weinshenker, Zhexing Wen, Stormi White, and Michael E. Zwick.

## Conflict of Interest

The authors have no competing financial interests related to this study to declare.

## Notes

### Competing Interest Statement

The authors have declared no competing interest.

