## Supplemental Methods and Results for "Convergent and distributed effects of the schizophrenia-associated 3q29 deletion on the human neural transcriptome"

### *Supplemental Information*

Table of contents:

**Extended Methods**……………..........................................................................................3-12
Overview…………………………………………………………………………………………3
Weighted gene co-expression network analysis (WGCNA)……………………………….……………4-6
*Samples used for network construction*………………………………………..….…..........4
*Data cleaning*……………………………..………………………………….…..….…...........4-5
*Network construction and module detection* .………………………………….….…..........5-6
Module preservation and quality analysis…..........................................................................7-8
Functional characterization of network modules harboring 3q29 genes…………………...…....8-9
Identification of meta-modules harboring 3q29 genes………......................................................9
Determination of module membership strength for 3q29 genes………………………….........9-10
Identification of prioritized driver genes…………………………....…………………......10-12
Functional characterization of prioritized driver genes………………….…………..….............12

**Extended Results**...................................................................................................................13-22
Unbiased gene co-expression network analysis reveals convergent and distributed effects of 3q29 interval genes across the adult human cortical transcriptome……………………..…….13-14
Pathway analysis points to functional involvement of the 3q29 locus in nervous-system functions and core aspects of cell biology……….…………………………………………..…….14-17
Network modules found to harbor 3q29 interval genes are robust and strongly reproducible in an independent test dataset………………………………………………………...........…….17-18
*UBXN7* is a highly connected cortical hub-gene predicted to play a crucial role in the neuropsychiatric sequela of 3q29Del…………………………………………………………...….18-19
Nine 3q29 interval genes form transcriptomic subnetworks enriched for known SZ, ASD and IDD-risk genes.……………………………………………..…………………………..…….....19-21
Disease-relevant driver genes prioritized by network analysis load onto key biological pathways linked to neuropsychiatric disorders.………..……………………………………...……21-22

**Availability of Data & Materials**.....................................................................................................23

**Supplemental References**……..........................................................................................24-29

**Supplemental Figures** (separate PDF file)
Fig. S1. Tissue sample attributes and donor phenotypes of the GTEx dataset used for reference network construction
Fig. S2. Pre-processing of the reference dataset: outlier removal
Fig. S3. Determination of the soft-thresholding power ($\beta$) in WGCNA
Fig. S4. Determination of network reproducibility and module preservation in an independent test dataset
Fig. S5. Individual module-preservation and quality statistics underlying composite Zsummary scores
Fig. S6. Publication numbers for 3q29 genes and historic schizophrenia spectrum disorder candidate genes

**Supplemental Tables** (separate multi-sheet Excel file)
Supplemental Tables 1 (separate multi-sheet Excel file)

Table S1.1. Sample attributes and donor phenotypes for test and reference networks.
Table S1.2. Network-derived gene-module pairs.
Table S1.3. Eigengene-based module connectivity measures (kME).
Table S1.4. Module preservation and quality statistics for network validation.
Table S1.5. Functional enrichment results for network modules harboring 3q29 interval genes.
Table S1.6. Cortical mRNA expression summaries of 3q29 interval genes obtained from the Human Protein Atlas.

Supplemental Tables 2 (separate multi-sheet Excel file)
Table S2.1. Network-derived top intramodular neighbors of 3q29 interval genes.
Table S2.2. Curated gene-sets for disease association analysis.
Table S2.3. Network-derived prioritized driver genes & hypergeometric test results assessing the overlap between curated gene-sets and top neighbors of 3q29 interval genes.
Table S2.4. Functional characterization of prioritized driver genes.

### Extended Methods

#### Overview:

For this analysis, we selected the most appropriate publicly available transcriptomic dataset and cleaned and processed the data before constructing an unsupervised gene co-expression network that can group functionally correlated genes into modules [1]. Genes participating in the same molecular and biological pathways tend to show correlated expression (co-expression) with each other, as they are expressed under the control of a coordinated transcriptional regulatory system [2]. Constructed on this principle, the transcriptomic network identified in this study amounts to a systems-level representation of gene-gene relationships in the healthy adult human prefrontal cortex (PFC), providing novel insights into network-level operations of understudied genes located in the recurrent 3q29 deletion (3q29Del) locus.

Given the strong genetic link between 3q29Del and risk for schizophrenia spectrum disorders (SZ; estimated odds ratio >40) [3], we placed our focus on revealing the co-regulation patterns of 3q29 interval genes as a function of their expression similarity during adulthood. This is a period when SZ typically manifests itself diagnostically, with peak onset in late adolescence (age 18-20 years) and early adulthood (age 20 to 40 years) [4], and a substantial proportion of patients becoming ill during middle adulthood (age 40 to 60 years) [5]. We also focused our analysis on uncovering gene-gene relationships in the PFC: a brain region that subserves a diverse range of cognitive and emotional operations, is most consistently implicated in the etiology of SZ [6] and may be particularly vulnerable to the effects of genetic disruption due to its protracted development [7]. To validate the reproducibility of this transcriptomic network, we used an independent test dataset that was demographically comparable to our reference dataset and assessed the preservation (i.e., how well-defined modules are in an independent test dataset) of each identified network module by permutation tests. Using a similar approach, we also assessed the quality (i.e., how well-defined modules are relative to the background) of each network module in repeated random splits of our reference dataset.

We next identified and interrogated the modules that were found to harbor at least one 3q29 interval gene for (1) functional enrichment, (2) higher-level organization into meta-modules, (3) highly connected "hub" genes and (4) "top" network neighbors of 3q29 interval genes. To generate testable hypotheses about which 3q29 interval genes may be causally linked to the major neuropsychiatric phenotypes of 3q29Del, we tested whether the top network neighbors of individual 3q29 interval genes show significant overrepresentation of known risk genes for SZ, or for two other clinical phenotypes associated with 3q29Del: autism spectrum disorders (ASD) and intellectual / developmental disability (IDD) ASD and IDD share genetic risk and pathogenic mechanisms with SZ, despite differences in the timing of clinical symptoms [8]. Based on the identified enrichment patterns, we developed a list of "prioritized driver genes", consisting of select 3q29 interval genes and their tightly correlated SZ, ASD and IDD-associated top network neighbors, which were subsequently found to converge onto canonical biological pathways.

We predict that dysregulation of these prioritized molecular targets, including a hub gene within the 3q29 interval, and their associated biological pathways may be implicated in the neuropsychiatric phenotype in 3q29Del syndrome. Overall, our findings highlight the advantage to using unbiased systems approaches that integrate gene-level information into a higher-order, network-level framework to infer gene function, particularly for understudied genes with disease relevance. The molecular pathways and the prioritized gene-set identified in this study will inform new directions of neurobiological inquiry that can mechanistically connect 3q29Del with severe mental illness. Our approach also highlights the study of normal function in non-pathological post-mortem tissue to further our understanding of psychiatric genetics, especially for rare genetic syndromes like 3q29Del, where access to post-mortem neural tissue from carriers is unavailable or limited. The steps described in this overview are described in great detail below.

**Weighted gene co-expression network analysis (WGCNA)**

*Samples used for network construction:*

To interrogate the functional consequences of the 3q29Del at the individual gene level, we employed unbiased weighted gene co-expression network analysis (WGCNA package [1, 9] in R, version 1.68) and organized the adult human cortical transcriptome into clusters (modules) of highly correlated genes (nodes). The network was constructed on publicly available RNA-Seq data obtained from the Genotype Tissue Expression Project (GTEx) [10] (Supp. Fig. S1), the largest multi-tissue open-data initiative using postmortem samples from human donors. 113 non-pathological post-mortem samples from the PFC (Brodmann Area 9) of male and female adults (age range = 20-79) with no known history of psychiatric or neurological disorder were included in this study. Transcriptome profiling was performed using Illumina TruSeq RNA sequencing as described in GTEx Consortium et al. (2013, 2015) [10]. The GTEx data (release version 6) used for the analyses described in this manuscript were downloaded from the GTEx portal (http://www.gtexportal.org/home/datasets/) on 01/08/2019; the corresponding dbGaP study accession number is phs000424.v6.p1.

*Data cleaning:*

Six outlier samples were removed from the original dataset prior to network construction to prevent a potential outlier-driven bias in network structure and module detection (Supp. Fig. S2). Inter-sample correlation (ISC) was used as an unbiased statistical diagnostic for identifying outliers with divergent gene expression profiles, defined as the Pearson's correlation between pairs of samples across the expression levels of all detected genes. Samples with a mean ISC greater than two standard deviations away from the mean of the sample-set were flagged as outliers and removed (as described in Oldham et al., 2008 [11]), reducing the sample size to 107. Details on sample attributes and donor phenotypes are available in Supp. Table S1.1 and Supp. Fig. S1.

Protein-coding genes were extracted from the dataset based on GENCODE v19 annotations for gene-type [12], followed by $\log_2$ transformation of gene expression values. For genes indexed by multiple splice variants, the data were further pre-processed to limit the transcriptome to a single transcript per gene. This step was conducted by using the *collapseRows* function of the WGCNA package in R (method = MaxMean), that identifies the transcript with fewest number of missing values and highest mean expression value across samples [13]. We limit the scope of this study to protein-coding genes with expression profiles summarized at the gene-level, in light of previously demonstrated drawbacks of isoform-level networks encompassing the non-coding transcriptome [14]. Inclusion of splice variants and non-coding RNAs increases network size by 100-fold [14], which in turn dramatically increases the computational resources necessary for network analysis. Although this high computational demand can theoretically be overcome by using a block-wise design where automatically selected subsets of the original data are used to build a co-expression network in a block-by-block fashion [1], previous work has shown that networks generated by block-wise design reflect only an approximation of networks generated by single-block design, which negatively influences the accuracy of subsequent module detection [14]. To avoid the shortcomings of a block-wise approach and pursue the most parsimonious hypothesis, our dataset included only protein-coding genes, with expression values summarized at the gene-level. This approach yielded a trimmed protein-coding gene expression matrix of 18,410 unique HGNC gene symbols.

To remove non-biological experimental variation, the dataset was adjusted for known batch effect (nucleic acid isolation batch) using the empirical Bayes framework employed by the *ComBat* function of the Surrogate Variable Analysis (SVA) package in R [15] (as described in [16]). The gene expression data were further corrected for age, sex, death classification (assessed by the 4-point Hardy Scale) and post-mortem interval (PMI)-mediated covariance [17], added to the regression model as categorical or continuous variables as applicable (Fig. 1b, Supp. Fig. S1). The residuals calculated from this model were carried to downstream analysis. Finally, since low-expressed or non-varying genes usually represent noise in the data, we used the *goodSamplesGenes* function of the WGCNA package in R to iteratively identify and remove genes and samples with greater than 50% missing entries (default parameter) and genes with zero variance. The normalized, outlier-removed, residualized cortical expression values of 18,410 protein-coding genes from 107 samples constitute the final dataset for construction of the network.

*Network construction and module detection:*

The single-block pipeline implemented in the WGCNA R package was employed to build a signed and weighted gene co-expression network using the final dataset. Co-expression similarity was defined as the biweight midcorrelation (bicor) coefficient between the expression profiles of gene-pairs calculated for all possible comparisons. Bicor is a median-based measure of co-expression that was chosen as a robust alternative to mean-based similarity metrics (i.e., Pearson correlation) in evaluating similarity in gene expression [18]. To capture the continuous nature of pairwise interactions in biological systems and accentuate strong positive correlations, the resulting co-expression similarity matrix was transformed into a signed and weighted adjacency matrix. This step was conducted by using the soft-thresholding procedure implemented in the *pickSoftThreshold* and *scaleFreePlot* functions of the WGCNA package in R. A soft-thresholding power ($\beta$) of 8 was identified as the lowest possible $\beta$ yielding a power-law degree distribution that approximately fits one with a scale-free network topology, (signed $R^2$ fit index = 0.8), while maintaining a relatively high mean connectivity (mean k > 100) (Fig. 1c, Supp. Fig. S3). The rationale behind our choice to establish a scale-free topology is the demonstrated biological relevance of its degree distribution as a unifying property of many biological networks in nature, where a few hub nodes have many connections while most nodes have few connections [19-22]. Additionally, note that a demonstrated advantage of weighted correlation networks is the high robustness of the network construction to the choice of $\beta$ parameter [23].

A signed adjacency matrix (Equation 1) was chosen over an unsigned adjacency matrix (Equation 2) to re-scale the underlying pairwise correlations from the [−1, 1] interval to the [0, 1] interval, as opposed to treating negative correlations as positive. In other words, since the adjacency matrix that underlies network construction is always non-negative, a signed network was chosen to respect the direction (up vs. down) of the co-expression relationships to prevent a clustering structure that mixes negatively correlated nodes, which often belong to different biological categories, with positively correlated nodes. The biological relevance of signed adjacency was demonstrated by previous work indicating that genes showing positive transcriptional correlation are more likely to exhibit known protein-protein interactions than uncorrelated or negatively correlated genes [11, 24, 25]. This approach is further supported by the recommendations of the creators of the WGCNA package in R (as described in https://peterlangfelder.com/2018/11/25/signed-or-unsigned-which-network-type-is-preferable/).

Signed adjacency $A_{ij}^{signed}$ for genes $i$ and $j$ is defined as:

$$A_{ij}^{signed} = \left|\frac{1+bicor\,(x_i,x_j)}{2}\right|^{\beta} \qquad (1)$$

Unsigned adjacency $A_{ij}^{unsigned}$ for genes $i$ and $j$ is defined as:

$$A_{ij}^{unsigned} = \left|bicor\,(x_i,x_j)\right|^{\beta} \qquad (2)$$

The resulting signed and weighted adjacency matrix was transformed into a topological overlap (TO) matrix to capture not only the correlation between pairs of genes but also the connections among "neighborhoods" of genes [26, 27]. The TO measure (TOM) between two genes is high if the genes have many overlapping network connections, yielding a network interconnectedness measure that is proportional to the number of common neighbors shared between a pair of nodes. Several studies have shown that gene/protein-pairs with higher TO are more likely to play a role in the same functional class than gene/protein-pairs with lower TO [27-32], demonstrating that TOM yields biologically meaningful modules that can successfully capture the co-expression profile of genes encoding interacting proteins. Additionally, TOM was shown by previous work to be more robust to identifying spurious connections than pairwise correlation alone [33].

To explore the clustering structure of the nodes underlying our undirected network, we conducted average linkage hierarchical clustering on pairwise TOM, following its transformation into a dissimilarity measure (dissTOM = 1-TOM). To define network modules, we adaptively pruned the branches of the resulting dendrogram by using the dynamic-hybrid-tree-cut algorithm of the WGCNA package in R (Fig. 1d). This method employs a bottom-up approach that considers the shape of dendrogram branches in identifying clusters, yielding improved detection of outlying cluster members. This method was chosen, since it has been shown to outperform the traditional fixed-height branch cutting method for identifying biologically meaningful clusters, particularly in networks that exhibit a nested dendrogram structure [34]. Standard parameters were used to conduct this analysis: cut height = 99% of the truncated height range in the dendrogram, module detection sensitivity (deep split) = 2, minimum module size = 30, signed network with partitioning about medoids (PAM) respecting the dendrogram. To avoid over-clustering, we set the smallest number of genes that can be considered a module (minimum module size) to 30; this is a standard parameter used in the literature to establish a compromise between large modules that are robust and biologically informative and small modules that are possibly informative but less robust [35-37]. This approach yielded 31 modules, with module-sizes ranging from 43 to 3,319 genes; similar ranges have been reported in other applications.

We summarized the gene expression profiles of individual modules by eigengenes [1], identified as the first principal component of the expression data in a given module. Construction of eigengenes amounts to a network-based data reduction method that serves as a means to effectively correlate entire modules. Leveraging this approach, we amalgamated modules with very similar expression profiles to eliminate spurious assignment of highly co-expressed genes into separate modules. To this end, we conducted average linkage hierarchical clustering of module eigengenes (ME) on a correlation-based dissimilarity metric (1-pairwise Pearson's correlation between MEs) and merged modules that are strongly correlated ($r > 0.8$ corresponding to cut height = 0.2) (Fig. 1d).

One of the resulting modules (grey module) was reserved for genes that could not be unequivocally assigned to any module (did not share similar co-expression patterns with the other genes of the network), as determined by the iterative refinement methodology used in our analysis pipeline to improve module detection. Therefore, the grey module was excluded from downstream analysis. Finally, we tested additional WGCNA parameters and determined that basic module structure in modules of interest (modules harboring 3q29 interval genes) was identifiable under variations of carefully selected algorithm parameters. A list of gene-sets for each identified module is provided in Supp. Table S1.2.

**Module preservation and quality analysis**

To validate the reproducibility of our identified modules, we evaluated network preservation in an independent transcriptomic dataset, hereafter referred to as the test dataset/network. Publicly available RNA-Seq data from the BrainSpan Developmental Transcriptome Project [38] was used to build the test network (Supp. Fig. S4). 30 non-pathological post-mortem samples from the PFC of male and female adults (age range = 18-37) with no known neurological or psychiatric disorder were used in this study. To exclude any confounding pathology, specimens had been confirmed by neuropathological evaluation to not contain obvious malformations, extensive neuronal loss, neuronal swelling, or dysmorphic neurons and neurites. Additionally, any donor with a reported prolonged agonal condition (i.e., coma, hypoxia, seizures etc.), ingestion of neurotoxic substances at the time of death, suicide, severe head injury, significant hemorrhages, prominent vascular abnormalities, tumors, prominent brain lesions, stroke, congenital neural abnormalities, and signs of neurodegeneration (i.e., amyloid plaques, Lewy bodies etc.) were also excluded (see technical white paper [39]). Transcriptome profiling was performed using Illumina TruSeq RNA sequencing. Details on sample attributes and donor phenotypes are available in Supp. Table S1.1 and Supp. Fig. S4. The BrainSpan data (Developmental Transcriptome Dataset; v10 summarized to genes) used for the analyses described in this manuscript were downloaded from the Allen Brain Atlas portal (https://www.brainspan.org/ static/download/) on 01/12/2019; the corresponding dbGaP study accession number is phs000755.v2.p1.

The test dataset was pre-processed following the same pipeline used for the GTEx reference dataset. We were unable to correct the normalized gene expression values of the test dataset for batch effect and death classification-mediated covariance, due to absence of the pertinent information from publicly available data. However, we note that the highly selective inclusion/exclusion criteria used by the BrainSpan project for tissue qualification [38] mitigates a potential confounding effect of death classification on gene expression. The inclusion of age, sex and PMI in the regression model was consistent with the pre-processing pipeline applied to the reference dataset. No outlier sample was identified by the above described ISC method. Notably, the test dataset was assembled by the aggregation of transcriptomic data from four sub-regions of the PFC (orbital, dorsolateral, ventrolateral, and medial PFC). To rule out a potential bias in test network construction driven by tissue-type, we conducted average linkage hierarchical clustering on the BrainSpan samples used in this study. The similarity metric for clustering was defined as the Pearson's correlation between gene expression levels of pairs of samples. The resulting dendrogram revealed no clustering pattern driven by tissue-type, ruling out PFC sub-region as a factor that could bias network construction (Supp. Fig. S4). Upon completion of pre-processing, the test dataset consisted of normalized and residualized gene expression values for 18,339 unique HGNC symbols from 30 samples.

To determine whether properties of within-module topology that were identified in our reference network were preserved in the test network, we calculated a composite network-based preservation statistic ($Z_{summary.pres}$) [40] for each module by using the *modulePreservation* function of the WGCNA package in R. $Z_{summary.pres}$ is a summary statistic that encompasses multiple density-based and connectivity-based preservation statistics that test 1) whether nodes sharing the same module in the reference network remain highly connected in the test network (i.e., are groups of genes defined as modules in the reference network denser than random groups of genes in the test network?) and 2) whether connectivity patterns between nodes underlying the reference network remain similar in the test network (i.e., do hub nodes of the reference network preserve their high degree of connectivity in the test network?). To determine whether the observed preservation statistics were higher than expected by chance and to derive a standardized Z score for each preservation statistic, we randomly permuted the module assignments in the test data (number of permutations = 200; twice as high as default) and derived a $Z_{summary.pres}$ score for each module. The resulting scores were evaluated according to established thresholds [40]: $Z_{summary.pres} < 2$ indicates no evidence for preservation, $2 < Z_{summary.pres} < 10$ indicates moderate evidence for preservation, and $Z_{summary.pres} > 10$ indicates strong evidence for preservation. To account for the dependence of the $Z_{summary.pres}$ score and permutation test p-values on module-size, we set the maximum module-size parameter of the *modulePreservation* function to 1000 genes (default parameter), reducing large modules by randomly sampling 1000 intra-modular nodes. Note that the specific goal of this preservation analysis was to determine the preservation strength of individual modules in relation to established $Z_{summary.pres}$ score thresholds, as opposed to comparing the preservation statistics of modules with different sizes to one another (i.e., determine whether module A is more preserved than module B). Hence, aggregating multiple preservation statistics into an informative $Z_{summary.pres}$ score constitutes a valid and advantageous approach for our purposes, despite its sensitivity to module-size.

In addition to measuring the density and connectivity-based preservation of each module between the reference and test networks, we measured the quality of the identified modules in the reference network without a reference to the test network. The goal of this analysis was to assess the robustness of the identified modules (i.e., how distinct is a given module from other modules in the reference network?) by calculating a composite quality statistic ($Z_{summary.qual}$) [40] for each module, as implemented in the *modulePreservation* function of the WGCNA package in R. This approach is akin to a cluster stability analysis [41] and employs a resampling technique that applies the module preservation statistics outlined above to repeated random splits of the reference data. The resulting $Z_{summary.qual}$ score indicates the robustness of a given module definition (hence the parameters selected for network construction and module detection) across networks created from the original reference data. The same $Z_{summary.pres}$ thresholds outlined above were used to evaluate the $Z_{summary.qual}$ scores. Finally, in line with the recommendations [40] of the creators of the WGCNA package, we also evaluated the individual preservation and quality statistics underlying the composite scores for each module (Supp. Table S1.4, Supp. Fig. S5). Results from the module preservation and quality analysis (Fig. 1e) revealed the replicable and robust nature of the network, thus we commenced interrogation of the network for insights into 3q29 interval gene function.

**Functional characterization of network modules harboring 3q29 genes**

Understanding the functions of the 21 protein-coding genes hemizygously deleted in 3q29Del syndrome is an obligatory step towards gaining insights into the cellular mechanisms underlying disease etiology. Others have shown that each gene of the human genome is estimated on average to be involved in ten biological functions [42]. Given that many of the genes in the 3q29Del interval are poorly annotated and some lack a functional annotation altogether (Supp. Fig. S6), we leveraged the transcriptomic co-expression approach to predict gene function. We screened each network module for membership of 3q29 interval genes and exploited major biological databases to derive a functional interpretation of the co-expression modules found to harbor at least one 3q29 interval gene (hereafter referred to as a 3q29 module).

Functional enrichment analyses of individual 3q29 modules (transformed into gene-sets) were run on the g:Profiler webserver (http://biit.cs.ut.ee/gprofiler; Ensembl version 96, Ensembl Genomes version 43) by using gene ontology biological processes (GO:BP), and Reactome (REACT) and Kyoto Encyclopedia of Genes and Genomes (KEGG) biological pathways. For high-confidence analysis, electronic GO annotations ("IEA"), which are assigned to gene products without curator verification (majority inferred *in silico*) and often cannot be traced to an experimental source, were discarded; filtering GO annotations based on evidence code has been recommended in previous literature to avoid erroneous results [43]. The statistical domain scope for functional enrichment analysis was set to only genes with at least one known annotation (default parameter) to establish an effective genomic background for statistical testing using the hypergeometric probability function. Enriched terms surpassing the adjusted g:SCS significance threshold of $P < 0.05$ were filtered for gene-set size (allowed range: 10-2,000 genes) and semantic similarity to improve the specificity and interpretability of our results [44]. The g:SCS method was our method of choice as it was shown to outperform standard approaches (i.e., Bonferroni correction, Benjamini-Hochberg False Discovery Rate) in estimating the true effect of multiple testing over the complex structure of GO that includes many overlapping functional categories [45]. The REVIGO webtool (http://revigo.irb.hr/) was used to reduce redundancy in the identified GO terms (default semantic similarity measure = SimRel, allowed similarity = 0.9 (large)) [46]. Top 10 biological pathways (REACT) found to be enriched in individual 3q29 modules were ranked by statistical significance level (adjusted $P < 0.05$) and are shown in the main results. Extended results are provided in Supp. Table S1.5.

Finally, findings of the functional enrichment analysis were also evaluated to determine whether identified co-expression modules reflect true biological signals as opposed to noise; this distinction was inferred by enrichment of the constituent genes for known biological processes and pathways. Our gene ontology-based approach complemented the module quality analysis described above.

#### Identification of meta-modules harboring 3q29 genes

Several studies have demonstrated that individual modules can be organized into biologically meaningful meta-modules that represent a higher-order organization of the transcriptome [47], likely reflecting pathway dependencies and synergistic function. To evaluate whether individual 3q29 modules clustered together within larger meta-modules of the co-expression network, we investigated the relationship among modules by leveraging their eigengenes (ME). We calculated the Pearson's correlations between all pairs of MEs and used this similarity metric to conduct average linkage hierarchical clustering of the MEs. Consistent with its use in other applications, we defined meta-modules as tight clusters of positively correlated MEs detectable as major branches of the resulting eigengene dendrogram [47]. The identified meta-modules were screened to identify the grouping patterns among 3q29 modules across the co-expression network.

This eigengene-based network reduction framework was further utilized to determine whether individual 3q29 modules that were found to partake in larger meta-modules were truly distinct from each other. To this end, gene ontology information was leveraged to identify common versus distinct enrichment of biological processes and pathways in individual 3q29 modules sharing a meta-module. This approach also complemented the module quality analysis described above.

#### Determination of module membership strength for 3q29 genes

To measure how strongly connected individual 3q29 genes are to their assigned modules, we used the *signedKME* function of the WGCNA package in R to calculate a module membership measure (kME) for each 3q29 gene. kME is an eigengene-based module connectivity measure, defined as the Pearson's correlation between the expression profile of a given gene and the eigengene (first principal component) of a given module[1]. An advantage of using kME as a network connectivity metric is that it allows the direct comparison of module membership values across modules that differ in size. This property distinguishes kME from degree-based measures of module connectivity, which are derived by summing a node's total number and strength of connections within a module and result in a metric that can be biased by module size.

A kME of 0 indicates that a gene is uncorrelated with an ME of interest and is thus unlikely to be a member of that module. In contrast, kME > 0.8 describes a hub gene that is highly connected to other genes in its module, hence predicted to be a crucial component of the overall function of that module. Previous work has shown that targeted disruption of a hub gene has a more deleterious effect on the ability of the network to function and leads to a larger number of phenotypic outcomes than the disruption of randomly selected genes or targeted deletion of less connected genes [48, 49]. These observations have led to the hypothesis that hub nodes of biological networks are typically associated with human disease genes. Indeed, several lines of evidence, particularly from cancer biology, have validated this hypothesis [50, 51]. One study has shown that topological features of disease-genes identified from OMIM's Morbid Map of the Human Genome disproportionately exhibit hub-gene characteristics, with protein products participating in more known protein-protein interactions than that of non-disease genes[52]

Note that there is no established definition of a hub node in network analysis, since the selection criteria can vary depending on the sparsity of the network; examples of hub gene definitions based on kME ≥ 0.7 exist in the literature. To generate rigorous WGCNA-based predictions, we adopted a conservative criterion that defines hub genes as nodes with kME > 0.8 ($P < 0.05$), a stringent threshold used by several other studies [53-55]. We annotated 3q29 interval genes that surpassed this kME threshold as hub genes, whose loss of function is predicted to produce a highly deleterious impact on the system.

Additionally, we screened kME values to identify 3q29 genes with very weak module membership. Given that kME quantifies how close a gene is to its assigned cluster of co-expressed genes, we excluded 3q29 genes with non-significant kMEs ($P > 0.05$) from downstream analysis. Notably, we limited our evaluations to intra-modular kME values that describe a gene's correlation with the eigengene of its assigned module. In rare instances, the kME of a given gene was found to be slightly higher for a module other than its assigned module. This finding stems from the fact that TOM yields a measure of network interconnectedness that is similar but not identical to an only correlation-based approach (as described by the creators of the WGCNA package at https://support.bioconductor.org/p/101579/). As noted previously, TOM-based similarity was shown to outperform correlation-based similarity in identifying biologically meaningful modules [33]. A list of kME values and corresponding p-values computed via correlation tests is provided in Supp. Table S1.3 for each node and module.

**Identification of prioritized driver genes**

3q29Del confers >40-fold increased risk for SZ and constitutes a shared risk factor for IDD and ASD. However, no single gene in this interval has been definitively associated with SZ, IDD, or ASD. To generate testable hypotheses about which 3q29 interval genes are causally linked to the major neuropsychiatric phenotypes associated with 3q29Del, we further leveraged our gene co-expression network. We adopted the guilt-by-association approach and screened the intra-modular subnetworks of individual 3q29 interval genes for a significant overlap with known SZ, ASD and IDD-risk genes. Guilt-by-association is a widely used principle that operates on the assumption that disease-associated genes are more closely connected to each other than random pairs of nodes in a network. Ample evidence demonstrates the high utility of the guilt-by-association approach in identifying novel disease risk genes [56-60].

Previous work has shown that testing the overrepresentation of known disease-risk genes in entire network modules harboring a large number of genes produces an inflated false positive rate[14]. To avoid this, we reduced 3q29 modules to top-neighbor-based subnetworks, maintaining only the close intra-modular connections of individual 3q29 genes. A top neighbor was defined as any node whose gene expression profile has a moderate-to-high correlation (Spearman's rho (ρ) ≥ 0.5) with a given 3q29 interval gene (considered a "seed" node) within the same module. Note that top neighbors were identified by a correlation-based hard-thresholding method applied only to intra-modular edges connecting seed-node pairs. Hence, the criterion for top neighbor identification combines the strength of scale free-topology and topological overlap principles of WGCNA for network construction and module detection, with subsequent application of a selective hard-thresholding method, resulting in a binary classification of top neighbors predicted to form direct functional links with 3q29 interval genes.

We conducted hypergeometric tests to determine whether top-neighbor-based intra-modular subnetworks of individual 3q29 genes are enriched for curated gene-sets with known SZ, ASD or IDD association. Six gene-sets were curated for this purpose from the following sources: 1) 93 IDD-risk genes enriched for damaging *de novo* mutations, identified by the Deciphering Developmental Disorders Study [61]; 2) 86 ASD-risk genes categorized by the Simons Foundation Autism Research Initiative (SFARI) [62, 63] as "high-confidence candidate risk genes" with strong evidence for ASD association (categories = 1 & 2; downloaded on 2/15/2019 from https://gene.sfari.org/database/); 3) 651 SZ-related genes shown by Meng et al. (2018) [64] to demonstrate significant differential expression in the postmortem brain tissue of SZ cases/controls, identified from the PsychENCODE BrainGVEX dataset [65]; 4) 636 SZ-related genes shown by Fromer et al. (2016) to demonstrate significant differential expression in the postmortem dorsolateral PFC tissue of SZ cases/controls, identified from the CommonMind Consortium dataset [66], 5) 340 SZ-risk genes adjacent to SZ-associated genetic loci, identified by the most recent genome-wide association study (GWAS) [67] conducted by the Psychiatric Genomics Consortium (PGC); 6) 290 SZ-risk genes with exonic *de novo* mutations, identified via the Neuropsychiatric Disorder *De Novo* Mutations Database [68] (downloaded on 2/17/2019 from http://www.wzgenomics.cn/NPdenovo/). Notably, these gene-sets were selectively curated from the literature to obtain a comprehensive yet reliable list of reported IDD, ASD and SZ-associated genetic variants spanning a wide range of the allele frequency spectrum. As noted previously, 3q29 genes with non-significant intra-modular kMEs ($P > 0.05$) (Fig. 2c) were excluded from this analysis.

To evaluate the specificity of the investigated disease enrichment patterns, we also tested the significance of the overlap between 3q29 subnetworks and negative control gene-sets associated with Parkinson's disease (PD), late-onset Alzheimer's disease (AD) and inflammatory bowel disease (IBD). The gene lists for these conditions were considered "negative controls," as they constitute disease phenotypes (two related and one unrelated to brain health) with no known association to 3q29Del. To rule out a potential bias that could be introduced to our enrichment analysis by differences in the sizes of curated gene-sets, common genetic variants associated with height (large gene-set size comparable to several SZ gene-set sizes) were included as another negative control. Four negative control gene-sets were curated for this purpose from the following sources: 1) 25 AD-risk genes identified by the largest published GWAS meta-analysis conducted by the Genetic and Environmental Risk in AD/Defining Genetic, Polygenic and Environmental Risk for AD Consortium (GERAD/PERADES) [69]; 2) 67 PD-risk genes identified by the largest published GWAS meta-analysis conducted by the International PD Genomics Consortium [70]; 3) 98 IBD-related genes identified by the International IBD Genetics Consortium as "strong positional candidate genes" in GWAS-identified risk loci [71]; 4) 479 height-associated genes identified by the largest published GWAS meta-analysis conducted by the Genetic Investigation of Anthropometric Traits Consortium (GIANT) [72].

To accurately measure the union-size of the possible matches between the curated gene-sets and WGCNA-derived 3q29 subnetworks, genes that could only be present in one set (non-protein-coding genes; gene-type annotated by GENCODE v19) were excluded from the overlap analysis, yielding the final gene-set sizes listed above. Notably, discrepancies in gene symbol alias usage were accounted for to ensure consistency in nomenclature. The background list for the hypergeometric test was determined as the total number of unique genes used to conduct WGCNA. The *GeneOverlap* package[73] in R (version 1.20.0) was used for this analysis. The hypergeometric p-values obtained by overlap analysis were corrected for multiple testing using the Benjamini-Hochberg procedure ($n$ = 10 gene-sets). 3q29 genes whose subnetworks were found to show a significant overlap with SZ, ASD and/or IDD risk genes (adjusted $P < 0.05$) were prioritized as driver genes, along with their SZ, ASD, and/or IDD-related top neighbors from the corresponding enriched disease gene-set. These prioritized driver genes are predicted to contribute to the emergence of the major neuropsychiatric phenotypes associated with 3q29Del. The list of curated gene-sets, WGCNA-derived top neighbors, WGCNA-derived prioritized driver genes and detailed results of the overlap analysis are provided in Supp. Tables S2.1-3.

### Functional characterization of prioritized driver genes

According to the “local hypothesis” proposed by the emerging paradigm of network medicine, the human transcriptome and proteome demonstrate non-random topological characteristics, where disease genes tend to interact with other disease genes and play distinct roles in disrupting the same biochemical process underlying a common pathophenotype [74]. Motivated by this theory, we sought to formulate testable hypotheses about the key biological mechanisms linking 3q29Del to SZ, ASD, and IDD by conducting a functional enrichment analysis on the union of our prioritized driver genes. We used the same analysis approach described above for testing the functional enrichment of 3q29 modules. The biological processes and pathways that were found to surpass the adjusted g:SCS significance threshold of p<0.05 were filtered for gene-set size and semantic similarity; the top 20 biological processes and pathways found to be enriched in our prioritized driver genes are shown in the main results. To provide a thorough illustration of all enriched GO:BP terms in the main results, GO:BP findings were further organized into a network visualization of related functional annotation categories. Extended results are provided in Supp. Table S2.5.

Overall, the transcriptomic network identified in this study is predicted to connect 3q29 interval genes with gene-sets outside the interval that participate in the same or overlapping biological process and associate with similar disease phenotypes. Perturbation of 3q29 interval gene dosage is expected to also perturb the functioning of network-partners outside the recurrent 3q29Del locus. However, note that the underlying structure of weighted gene co-expression networks is agnostic to the mechanistic order of cellular and molecular events. The information necessary to derive the order of biological interactions is not an explicit outcome of gene co-expression itself, since such inferences require time-dependent analysis of combinatorial interactions between nodes. As a result, some of the network partners identified in this study are expected to function upstream of their co-expressed 3q29 interval partner, and likely not be affected by 3q29Del.

### Extended Results

#### Unbiased gene co-expression network analysis reveals convergent and distributed effects of 3q29 interval genes across the adult human cortical transcriptome.

Our WGCNA-based unsupervised network analysis approach, applied to publicly available high-throughput GTEx data, revealed that the protein-coding transcriptome of the healthy adult human PFC can be organized into a co-expression network of 19 modules (labeled by color) (Fig. 1d, Fig. 2a). The identified modules group genes with highly similar expression profiles into densely interconnected clusters, which likely represent shared function and co-regulation. One of the identified modules (the grey module) contained genes that could not be unequivocally assigned to any module; thus, it was excluded from downstream analysis. The resulting module sizes (number of genes assigned to a module) ranged from 43 (steel blue) to 4,746 (green) genes, with an average module size of 1,014 genes (excluding grey module). Similar ranges have been reported in other network analysis applications. A list of gene-sets for each identified module is provided in Supp. Table S1.2.

The 21 protein-coding genes located in the 3q29 interval were found to cluster into seven network modules, which represent local regions of the human protein-coding genome that demonstrate coordinated expression with the 3q29 locus (Fig. 2a). These modules are referred to as 3q29 modules, and were labeled as black (size = 1,170 genes), brown (size = 1,972 genes), dark turquoise (size = 496 genes), green (size = 4,746 genes), magenta (size = 1,437 genes), midnight blue (size = 1,414 genes), and turquoise (size = 3,319 genes) modules. Moreover, 18 / 21 3q29 interval genes were found to concentrate into just four modules (brown, green, midnight blue, turquoise) (Fig. 2a), suggesting that the haploinsufficiency of the 3q29 locus may perturb the same biological processes via multiple hits, cumulatively disrupting redundancy and compensatory resiliency in the normative regulation of cellular functions. Simultaneously, leading candidate genes *DLG1* (black) and *PAK2* (dark turquoise), which were previously hypothesized to contribute to neuropsychiatric pathology, were found in opposite branches of the network, demonstrating the potential distributed effects of this CNV across the transcriptomic landscape (Fig. 2a). The modular allocation of 3q29 interval genes is listed below:

| ***Modules harboring 3q29 interval genes ("3q29 Modules")*** | | |
|---|---|---|
| **Network module** | **Number of 3q29 interval genes** | **Constituent 3q29 interval gene symbols** |
| Black | 1 | *DLG1* |
| Brown | 4 | *NCBP2, TFRC, TM4SF19, ZDHHC19* |
| Dark turquoise | 1 | *PAK2* |
| Green | 6 | *BDH1, PIGZ, PCYT1A, SMCO1, SLC51A, MFI2* |
| Magenta | 1 | *RNF168* |
| Midnight blue | 3 | *UBXN7, SENP5, WDR53* |
| Turquoise | 5 | *CEP19, FBXO45, PIGX, TCTEX1D2, NRROS* |

The average linkage hierarchical clustering of the module eigengenes revealed that the identified modules further clustered into three higher level meta-modules (clusters of highly correlated modules), detected as major branches of the resulting eigengene dendrogram (Fig. 2a, 2b). The midnight blue and black 3q29 modules were found to cluster together within the first meta-module, grouping three 3q29 interval genes: *DLG1, UBXN7, SENP5* and *WDR53*. The magenta, green and turquoise 3q29 modules clustered together within the second meta-module, grouping 12 3q29 interval genes: *RNF168, BDH1, PIGZ, PCYT1A, SMCO1, SLC51A, MFI2, CEP19, FBXO45, PIGX, TCTEX1D2* and *NRROS.* Finally, the brown and dark turquoise 3q29 modules clustered together within the third meta-module, grouping five 3q29 interval genes: *NCBP2, TFRC, TM4SF19, ZDHHC19* and *PAK2.* The observed grouping, as well as the segregation, of sets of 3q29 modules into distinct meta-modules represents a higher-order transcriptomic organization of the 3q29 locus, which likely reflects pathway dependencies and interactions between biological processes involving 3q29 interval genes. These findings suggest that, rather than functioning as independent non-interacting units, sets of 3q29 interval genes and their co-expressed network partners may work in synergy at both the module and meta-module levels of transcriptomic organization (Fig. 2a, 2b), and likely constitute interacting sources of pathology in 3q29Del syndrome.

**Pathway analysis points to functional involvement of the 3q29 locus in nervous-system functions and core aspects of cell biology.**

The functional enrichment analysis of individual 3q29 modules showed that the constituent genes of each module load highly onto canonical biological processes and pathways (Fig. 1E). These functional enrichment findings validate that our co-expression-based 3q29 modules reflect clustering dynamics that are biologically meaningful.

Functional characterization of the black module: We observed that the biological processes and pathways that were significantly overrepresented in the black module mainly encompass terms related to regulation of gene expression and maintenance of the integrity of the cellular genome. Based on the ranking of p-values adjusted for multiple testing, the top biological pathways (annotated by the Reactome database) that were overrepresented in the black module include metabolism of RNA (REAC:R-HSA-8953854, adjusted $P$ = 2.45E-07), processing of capped intron-containing pre-mRNA (REAC:R-HSA-72203, adjusted $P$ = 7.74E-04), chromatin organization (REAC:R-HSA-4839726, adjusted $P$ = 6.36E-03), mRNA splicing (REAC:R-HSA-72172, adjusted $P$ = 2.11E-02), post-translational protein modification (REAC:R-HSA-597592, adjusted $P$ = 2.33E-02), DNA Repair (REAC:R-HSA-73894, adjusted $P$ = 4.02E-02) and tRNA processing (REAC:R-HSA-72306, adjusted $P$ = 4.97E-02).

Functional characterization of the midnight-blue module: Similarly, the midnight-blue module, which is in the same meta-module as the black module, was found to be enriched for biological pathways and processes that are involved in DNA repair and regulation of gene expression at the levels of transcription and translation, as well as cellular response to stress. An important functional signature that set the midnight blue module apart from the black module was its specific enrichment for terms related to cell cycle regulation. The top biological pathways that were overrepresented in the midnight blue module include gene expression (transcription) (REAC:R-HSA-74160, adjusted $P$ = 2.10E-39), metabolism of RNA (REAC:R-HSA-8953854, adjusted $P$ = 1.64E-06), DNA double-strand break repair (REAC:R-HSA-5693532, adjusted $P$ = 9.79E-04), mRNA 3'-end processing (REAC:R-HSA-72187, adjusted $P$ = 5.49E-04), cell cycle (REAC:R-HSA-1640170, adjusted $P$ = 3.26E-06), mRNA splicing (major pathway) (REAC:R-HSA-72163, adjusted $P$ = 5.33E-03), cell cycle checkpoints (REAC:R-HSA-69620, adjusted $P$ = 5.73E-03), transport of mature mRNA derived from an intron-containing transcript (REAC:R-HSA-159236, adjusted $P$ = 1.53E-02), and transcriptional regulation by the tumor suppressor TP53 (REAC:R-HSA-3700989, adjusted $P$ = 2.10E-02).

Functional characterization of the brown module: Assessment of shared function among constituent genes of the brown module revealed primary enrichment for biological pathways and processes involved in cellular metabolism and mitochondrial function. The top biological pathways that were overrepresented in the brown module include fatty acid metabolism (REAC:R-HSA-8978868, adjusted $P$ = 7.59E-08), mitochondrial fatty acid beta-oxidation (REAC:R-HSA-77289, adjusted *P* = 8.33E-05), chondroitin sulfate/dermatan sulfate metabolism (REAC:R-HSA-1793185, adjusted *P* = 1.42E-03), peroxisomal protein import (REAC:R-HSA-9033241, adjusted *P* = 1.79E-03), metabolism of fat-soluble vitamins (REAC:R-HSA-6806667, adjusted *P* = 4.99E-03), peptide hormone biosynthesis (REAC:R-HSA-209952, adjusted *P* = 8.72E-03), solute-carrier (SLC)-mediated transmembrane transport (REAC:R-HSA-425407, adjusted *P* = 8.94E-03), and diseases associated with glycosaminoglycan metabolism (REAC:R-HSA-3560782, adjusted *P* = 1.30E-02). Notably, the brown module was also found to be enriched for two canonical KEGG-annotated pathways: the Hippo signaling pathway (KEGG:04390, adjusted *P* = 4.82E-03) and the Wnt signaling pathway (KEGG:04310, adjusted *P* = 4.96E-02), which play crucial roles in growth and developmental pathways with substantial cross-talk [75].

Functional characterization of the dark turquoise module: The dark turquoise module was found to coalesce genes that are enriched for biological functions in epigenetic regulation of gene expression, as well as in signal transduction pathways that are mediated by Rho GTPases. This latter function is at least in part attributable to *PAK2*, the only 3q29 interval gene assigned to this module, which encodes a known Rho GTPase effector. Intriguingly, this module was also found to be enriched for a functional role in estrogen receptor-mediated signaling. In particular, the top terms that were overrepresented in the dark turquoise module include estrogen-dependent gene expression (REAC:R-HSA-9018519, adjusted *P* = 5.08E-06), estrogen receptor (ESR)-mediated signaling (REAC:R-HSA-8939211, adjusted *P* = 9.52E-06), Rho GTPase effectors (REAC:R-HSA-195258, adjusted *P* = 1.35E-05), SIRT1 negatively regulates rRNA expression (REAC:R-HSA-427359, adjusted *P* = 2.09E-05), gene silencing by RNA (REAC:R-HSA-211000, adjusted *P* = 3.95E-05), nucleosome assembly (REAC:R-HSA-774815, adjusted *P* = 4.75E-05), chromatin modifying enzymes (REAC:R-HSA-3247509, adjusted *P* = 5.12E-05), signaling by nuclear receptors (REAC:R-HSA-9006931, adjusted *P* = 8.46E-05), meiotic synapsis (REAC:R-HSA-1221632, adjusted *P* = 8.73E-05), and epigenetic regulation of gene expression (REAC:R-HSA-212165, adjusted *P* = 1.44E-04). Notably, the dark turquoise module was found to share the same meta-module as the metabolism-related brown module, suggesting the involvement of several 3q29 interval genes in a hierarchical transcriptomic control structure that interconnects Rho GTPase-mediated signaling cascades and estrogen-regulated signal transduction pathways with metabolic regulation. Emerging empirical findings demonstrate the existence of a crosstalk between these fundamental pathways [76-78], supporting the biological relevance of the co-expression-based clustering patterns underlying the meta-module that harbors the brown and dark turquoise modules identified in this study.

Functional characterization of the turquoise module: The biological processes that were overrepresented in the turquoise module primarily encompass nervous-system specific terms comprising the regulation of nervous system development and function. Other enriched biological functions that are non-specific but cardinal to nervous-system operations involve ion transport, calcium signaling, cyclic adenosine monophosphate (cAMP)-dependent signal transduction and cell projection organization. Specifically, the top Reactome-based biological pathways that were enriched in the turquoise module include neuronal system (REAC:R-HSA-112316, adjusted *P* = 2.98E-12), protein-protein interactions at synapses (REAC:R-HSA-6794362, adjusted *P* = 7.12E-07), neurexins and neuroligins (REAC:R-HSA-6794361, adjusted *P* = 3.58E-06), transmission across chemical synapses (REAC:R-HSA-112315, adjusted *P* = 6.31E-06), neurotransmitter receptors and postsynaptic signal transmission (REAC:R-HSA-112314, adjusted *P* = 9.47E-05), serotonin receptors (REAC:R-HSA-390666, adjusted *P* = 6.42E-03), the citric acid (TCA) cycle and respiratory electron transport (REAC:R-HSA-1428517, adjusted *P* = 7.00E-03), and unblocking of N-methyl-D-aspartate (NMDA) receptors, glutamate binding and activation (REAC:R-HAS 438066, adjusted *P* = 1.41E-02). Complementing these biological pathways, GO biological processes that were found to be enriched in the turquoise module include regulation of synaptic plasticity (GO:0048167, adjusted *P* = 6.68E-03), cognition (GO:0050890, adjusted *P* = 2.91E-02), neuron differentiation (GO:0030182, adjusted *P* = 3.19E-02), long-term potentiation (GO:0060291, adjusted *P* = 3.81E-02) and learning and memory (GO:0007611, adjusted *P* = 4.20E-02). Dysregulation of these biological processes and pathways has been implicated in the etiology of major neuropsychiatric and neurodevelopmental disorders, including SZ and ASD [79, 80]. Hence, the observed functional enrichment profile highlights a likely pivotal role for the coordinated expression of the 3q29 interval genes and network partners that cluster in the turquoise module in establishing and maintaining the healthy functioning of the brain.

<u>Functional characterization of the green module:</u> Similar to the turquoise module, the pathway enrichment analysis of the genes constituting the green module revealed primary enrichment for shared function in several nervous-system specific biological processes. The overarching functional characteristics of this module are regulation of nervous system development, interactions between neuroactive ligands and receptors, synaptic vesicle cycle, intracellular trafficking systems (i.e., vesicle-mediated synaptic transport) and synapse assembly. The top Reactome-annotated biological pathways that were found to be enriched in the turquoise module include neuronal system (REAC:R-HSA-112316, adjusted *P* = 7.76E-07), the role of GTSE1 in G2/M progression after G2 checkpoint (REAC:R-HSA-8852276, adjusted *P* = 6.14E-04), potassium channels (REAC:R-HSA-1296071, adjusted *P* = 2.99E-03), L1 cell adhesion molecule (L1CAM) interactions (REAC:R-HSA-373760, adjusted *P* = 8.69E-03), transmission across chemical synapses (REAC:R-HSA-112315, adjusted *P* = 1.66E-02), G protein-coupled receptor (GPCR) ligand binding (REAC:R-HSA-500792, adjusted *P* = 1.74E-02), G alpha (q) signaling events (REAC:R-HSA-416476, adjusted *P* = 2.37E-02), recycling pathway of L1 (REAC:R-HSA-437239, adjusted *P* = 3.93E-02), coat protein complex I (COPI)-mediated anterograde transport (REAC:R-HSA-6807878, adjusted *P* = 4.27E-02), and adenosine triphosphate-binding cassette (ABC) transporter disorders (REAC:R-HSA-5619084, adjusted *P* = 4.36E-02). The observed nervous system-specific functional enrichment findings suggest heightened disease-relevance for the 3q29 interval genes and intra-modular partners that coalesce in the green module. The neuropathology-associated functional characterization of the green module parallels that of the turquoise module, which shares the same meta-module as the green module. This functional overlap, which was identified agnostically to meta-module membership, presents further support for the utility of our approach in detecting biologically meaningful non-random network structures that organize gene expression in the healthy adult human cortex.

<u>Functional characterization of the magenta module:</u> The magenta module was found to be predominantly enriched for biological processes and pathways involved in post-translational protein modifications by small protein conjugation or removal, ubiquitin-dependent protein catabolism, intracellular protein transport and localization, and the ubiquitin-proteasome system. Hence, coordinated expression of the 3q29 interval genes and network partners that participate in the magenta module likely plays an important role in controlling the modification and spatiotemporal colocalization of substrates necessary for a variety of intracellular interactions. Moreover, pathway enrichment analysis revealed a link between magenta module genes and the initiation of MHC class I (MHC-I)-dependent immune responses, driven by a genomic locus that is increasingly implicated in the etiology of SZ [81]. MHC-I antigen presentation has been shown to strictly depend on peptide supply by the ubiquitin-proteasome system to initiate an effective adaptive immune response[82]; thus, the simultaneous enrichment of these interacting processes in a single module supports the biological relevance of the identified pattern of clustering. Specifically, the top Reactome-annotated biological pathways that were found to be significantly overrepresented in the magenta module include post-translational protein modification (REAC:R-HSA-597592, adjusted *P* = 1.22E-07), gene expression (transcription) (REAC:R-HSA-74160, adjusted *P* = 7.98E-06), MHC-I mediated antigen processing and presentation (REAC:R-HSA-

983169, adjusted $P$ = 1.69E-05), antigen processing: ubiquitination and proteasome degradation (REAC:R-HSA-983168, adjusted $P$ = 2.78E-05), RNA polymerase II transcription (REAC:R-HSA-73857, adjusted $P$ = 1.05E-04), signaling by TGF-beta family members (REAC:R-HSA-9006936, adjusted $P$ = 2.87E-03), protein ubiquitination (REAC:R-HSA-8852135, adjusted $P$ = 1.14E-02), and sumoylation (REAC:R-HSA-2990846, adjusted $P$ = 1.94E-02). Overall, the functional profile of the magenta module encompasses many known regulators of brain function, including synapse formation and trans-synaptic signaling. Hence, the functional characteristics of the magenta module complement that of its meta-module partners, the green and turquoise modules.

Taken together, functional characterization of the 3q29 modules (Fig. 2d) point to novel mechanisms of shared or overlapping action for sets of 3q29 interval genes that cluster in the same network module and further coalesce in the same meta-module. Simultaneously, the variety of biological pathways that were found to be enriched in 3q29 modules suggests distributed involvement of this locus in not only nervous-system specific functions, such as regulation and organization of synaptic signaling and components, but also in core aspects of cell biology, including cellular metabolism, transcriptional regulation, protein modifications, and cell cycle regulation. Extended results of the pathway enrichment analysis are provided in Supp. Table S1.5.

**Network modules found to harbor 3q29 interval genes are robust and strongly reproducible in an independent test dataset.**

To ensure the reproducibility of our network analysis results, we tested the preservation of various properties of graph structure that underly the modules identified in this study with respect to an independent dataset obtained from the BrainSpan Project (Supp. Fig. S4). We calculated multiple density-based and connectivity-based preservation statistics for each module using a permutation test procedure and summarized the observed statistics by a composite Z-statistic, $Z_{summary.pres}$. All identified modules, except for the grey module (unassigned genes), were found to be successfully preserved in the test network ($Z_{summary.pres}$ > 2) (Fig. 1e). Specifically, 3/18 modules exhibited moderate evidence of preservation (2 < $Z_{summary.pres}$ < 10), and 15/18 modules, including all 3q29 modules, exhibited strong evidence of preservation ($Z_{summary.pres}$ > 10) (Fig. 1e). Moreover, the resulting composite preservation statistics of all 3q29 modules were substantially higher than that of a randomly drawn sample of 1000 genes that represent the entire reference network as a single artificial module (labeled as the gold module, $Z_{summary.pres.gold}$ = 6.78).

In addition to preservation statistics, we calculated multiple module quality statistics that measure how well-defined or robust the boundaries of individual modules are in the reference network. By employing a resampling technique that applies module preservation statistics to repeated random splits of our reference data, we obtained a composite Z-statistic for each module ($Z_{summary.qual}$) that standardizes and summarizes multiple cluster quality statistics. All 18 modules showed strong evidence for high cluster quality ($Z_{summary.qual}$ > 10), revealing robust module definitions (Supp. Fig. S6C). Specifically, all 3q29 modules had a $Z_{summary.qual}$ score ≥ 20 (Fig. 1e).

Finally, in line with the recommendations of the creators of the WGCNA package[40], we also evaluated the individual preservation and quality statistics underlying the composite $Z_{summary.pres}$ and $Z_{summary.qual}$ scores derived for each module. Individual module preservation statistics mostly converge on the finding that (1) nodes sharing the same module in the reference network remain highly connected in the test network and (2) connectivity patterns between nodes underlying the reference network remain similar in the test network (Supp. Fig. S5). Similarly, individual module quality statistics predominantly indicate strong evidence for high cluster quality in all identified modules across networks created from random splits of the original reference data (Supp. Fig. S5).

Overall, these findings support the strong reproducibility and robustness of our 3q29 modules, allowing high-confidence screening of the transcriptomic connectivity patterns formed by 3q29 interval genes in the healthy adult human PFC. See Supp. Table S1.4 for detailed results of the module preservation and quality analysis.

**UBXN7 is a highly connected cortical hub-gene predicted to play a crucial role in the neuropsychiatric sequela of 3q29Del.**

To measure how strongly connected individual 3q29 interval genes are to their assigned network modules, we calculated the eigengene-based module connectivity measure (kME) of each 3q29 interval gene for its respective module (Fig. 2c). To reiterate, this measure quantifies how close a node is to its assigned module and can be applied to identify hub genes (kME > 0.8, $P < 0.05$), which are highly correlated with their module eigengene and exhibit high connectivity in their module. Intriguingly, our results revealed that *UBXN7*, an understudied and poorly annotated 3q29 interval gene, is a hub gene of its module (kME = 0.84, $P$ = 8.33E-30, midnight blue module size = 1,414 genes). Topological features of known disease-genes have been shown to disproportionately exhibit hub-gene characteristics compared to non-disease genes [52]. Supported by this literature, we predict that (1) *UBXN7* exerts central influence on a large network of co-expressed genes, and (2) loss of function mutations in *UBXN7* can cause major dysfunction in the biological pathways involving this gene. Consequently, we prioritize *UBXN7* as a major driver gene with likely disease relevance in 3q29Del. Intra-modular kME values of individual 3q29 genes, ranked from highest to lowest, are listed below:

| ***Eigengene-based module connectivity strength of 3q29 interval genes*** | | | |
|---|---|---|---|
| **3q29 interval gene** | **Network module** | **Intra-modular kME** | **Associated p-value** |
| *UBXN7* | Midnight blue | 0.84 | 8.33E-30 |
| *SENP5* | Midnight blue | 0.74 | 8.62E-20 |
| *PAK2* | Dark turquoise | 0.74 | 9.77E-20 |
| *PIGX* | Turquoise | 0.71 | 1.81E-17 |
| *CEP19* | Turquoise | 0.70 | 3.13E-17 |
| *RNF168* | Magenta | 0.70 | 5.06E-17 |
| *NCBP2* | Brown | 0.68 | 8.78E-16 |
| *PIGZ* | Green | 0.68 | 1.08E-15 |
| *BDH1* | Green | 0.64 | 8.32E-14 |
| *FBXO45* | Turquoise | 0.62 | 1.46E-12 |
| *WDR53* | Midnight blue | 0.59 | 2.04E-11 |
| *DLG1* | Black | 0.56 | 4.38E-10 |
| *TCTEX1D2* | Turquoise | 0.51 | 2.62E-08 |
| *NRROS* | Turquoise | 0.43 | 4.52E-06 |
| *PCYT1A* | Green | 0.40 | 2.00E-05 |
| *TFRC* | Brown | 0.39 | 3.56E-05 |
| *ZDHHC19* | Brown | 0.30 | 1.57E-03 |
| *TM4SF19* | Brown | 0.26 | 7.39E-03 |

| | | | |
|---|---|---|---|
| *SLC51A* | Green | 0.17 | 0.09 |
| *SMCO1* | Green | 0.11 | 0.25 |
| *MFI2* | Green | 0.09 | 0.35 |

Moreover, evaluation of the module membership strengths of 3q29 interval genes revealed that *SMCO1 (*kME = 0.11, *P* = 0.25*), SLC51A (*kME = 0.17, *P* = 0.09*) and MFI2 (*kME = 0.09, *P* = 0.35*)* have non-significant kMEs for their assigned module, suggesting poor module connectivity. The mRNA expression summaries obtained from the Human Protein Atlas [83] (HPA) for 3q29 interval genes indicate nearly negligible or very low mRNA expression levels for *SMCO1* (consensus normalized expression value = 0.1), *SLC51A,* (consensus normalized expression value = 0.4), *and MFI2* (consensus normalized expression value = 2.8) in the human cerebral cortex. These data indicate the low abundance of these 3q29 interval genes in our tissue of interest, which likely relates to their peripheral network assignments in our analysis. Consequently, *SMCO1, SLC51A* and *MFI2* were excluded from downstream analysis to ensure accurate refinement of tight network connections formed by 3q29 interval genes.

A list of kME values and associated p-values for all network node-module pairs is provided in Supp. Table S1.3. A list of mRNA expression summaries obtained from the HPA for all 3q29 interval genes is provided in Supp. Table S1.6.

**Nine 3q29 interval genes form transcriptomic subnetworks enriched for known SZ, ASD and IDD-risk genes.**

To systematically generate testable hypotheses regarding which 3q29 interval genes are causally linked to the major neuropsychiatric phenotypes associated with 3q29Del, we reduced 3q29 modules to strongly connected top neighbors of individual 3q29 genes and screened the resulting top neighbors for a significant overlap with known SZ, ASD or IDD-risk genes. To reiterate, this approach leverages the extensively validated principle of guilt-by-association, which postulates that the disease-relevance of a particular gene is partially a property determined by its relationships in a biological network.

A top neighbor was defined as any node whose gene expression profile has a moderate-to-high pairwise correlation ($\rho \geq 0.5$, $P < 0.05$) with a 3q29 interval gene within the same module. Intriguingly, our results revealed that several 3q29 interval genes are among the top neighbors of one another within the same module. *FBXO45* ($\rho = 0.5$, *P* = 5.43E-09) and *PIGX* ($\rho = 0.6$, *P* = 1.24E-10) were identified as top-neighbors of *CEP19* in the turquoise module. Similarly, *SENP5* and *WDR53* were top-neighbors of each other ($\rho = 0.5$, *P* = 1.05E-07) in the midnight blue module. This finding further suggests that the correlated activity of subsets of 3q29 interval genes may converge upon the same or synchronized multicomponent biological processes in the adult PFC.

Moreover, *TM4SF19 (0 top neighbors)* and *ZDHHC19 (3 top neighbors)* were found to have no or < 5 intra-modular partners in the brown module that met the correlation threshold to qualify as top neighbors. Similar to *SMCO1, SLC51A* and *MFI2,* the mRNA expression summaries obtained from the HPA[83] for *TM4SF19* (consensus normalized expression value = 0.5) and *ZDHHC19* (consensus normalized expression value = 0) indicate negligible or very low mRNA expression levels in the human cerebral cortex. These data independently indicate the low abundance of these 3q29 interval genes in our tissue of interest, which likely reflects the reason behind their lack of strongly connected top neighbors in our network analysis. Hence, *TM4SF19* and *ZDHHC19* were excluded from our downstream disease-association analysis, along with *SMCO1, SLC51A* and *MFI2,* which were deprioritized earlier due to poor module connectivity.

The intra-modular top neighbors of the remaining 16 3q29 interval genes were interrogated for overlap with six curated lists of evidence-based IDD, ASD or SZ-risk genes, spanning a wide range of the allele frequency spectrum (Fig. 3a). Hypergeometric test results, corrected for multiple testing, revealed a significant overrepresentation of one or more of these established risk gene-sets among the top neighbors of nine 3q29 interval genes (adjusted $P$ < 0.05): *BDH1, CEP19*, *DLG1*, *FBXO45, PIGZ, RNF168, SENP5, UBXN7* and *WDR53* (Fig. 3b). Details of the enrichment results with respect to module membership are provided below.

In the black module, top neighbors of *DLG1* (gene-set size = 294) were found to be enriched for known SZ-risk genes from the exonic *de novo* mutations gene-set (adjusted $P$ = 1.11E-05). This identified intersection comprises 18 unique top neighbors of *DLG1* with known SZ association. Note that *DLG1* was the only 3q29 interval gene that clustered in the black module.

In the midnight blue module, all three constituent 3q29 interval genes had top neighbors that loaded highly onto known IDD, ASD and/or SZ-risk genes. Particularly, top neighbors of *UBXN7* (gene-set size = 811) were enriched for known IDD (overlap size = 14, adjusted $P$ = 3.05E-04), ASD (overlap size = 15, adjusted $P$ = 5.46E-05), and SZ-risk genes from the CommonMind case-control gene-set (overlap size = 45, adjusted P = 4.20E-03). In addition, top neighbors of *SENP5* (gene-set size = 713) were enriched for known IDD (overlap size = 11, adjusted $P$ = 5.05E-03) and ASD-risk genes (overlap size = 12, adjusted $P$ = 1.28E-03). Similarly, top neighbors of *WDR53* (gene-set size = 278) had a significant overlap with known IDD (overlap size = 6, adjusted $P$ = 1.51E-02) and ASD-risk genes (overlap size = 6, adjusted $P$ = 1.51E-02). The union of these identified intersections adds up to a total of 67 unique top neighbors with known IDD, ASD and/or SZ association in this module.

In the green module, only two out of the six constituent 3q29 interval genes were found to have top neighbors that showed a significant overrepresentation of SZ-risk genes. Specifically, top neighbors of *BDH1* (gene-set size = 1008) were enriched for known SZ-risk genes from the CommonMind case-control gene-set (overlap size = 66, adjusted $P$ = 4.49E-06). Similarly, top neighbors of *PIGZ* (gene-set size = 995) were also enriched for known SZ-risk genes from the CommonMind case-control gene-set (overlap size = 58, adjusted $P$ = 7.03E-04). The union of these identified intersections adds up to a total of 75 unique top neighbors with known SZ association in this module.

In the magenta module, top neighbors of *RNF168* (gene-set size = 556) had a significant overlap with known IDD (overlap size = 9, adjusted $P$ = 1.07E-02) and SZ-risk genes from the CommonMind case-control gene-set (overlap size = 65, adjusted $P$ = 5.21E-17). The union of these identified intersections adds up to a total of 73 unique top neighbors with known SZ and/or IDD association in this module. Note that *RNF168* was the only 3q29 interval gene that clustered in the magenta module.

In the turquoise module, only two out of the five constituent 3q29 interval genes had top neighbors that loaded highly onto known IDD, ASD and/or SZ-risk genes. Specifically, top neighbors of *CEP19* (gene-set size = 1161) were enriched for known IDD (overlap size = 16 , adjusted $P$ = 1.64E-03), ASD (overlap size = 15, adjusted $P$ = 1.64E-03), and SZ-risk genes from the exonic *de novo* mutations gene-set (overlap size = 29, adjusted P = 3.32E-02). In addition, top neighbors of *FBXO45* (gene-set size = 1101) were also enriched for known IDD-risk genes (overlap size = 14, adjusted $P$ = 1.35E-02). The union of these identified intersections adds up to a total of 51 unique top neighbors with known IDD and/or SZ association in this module.

Finally, there was no statistically significant evidence for overrepresentation of known disease genes of interest among the top neighbors of 3q29 interval genes that clustered in the brown or dark turquoise modules.

To evaluate the specificity of the identified disease enrichment patterns, we also tested the top neighbors of 16 3q29 interval genes for overlap with known PD, late-onset AD and IBD risk genes. These disease phenotypes have no known link to 3q29Del syndrome, thus, genetic risk loci associated with these conditions were considered negative controls in this network analysis. In addition, a large list of common variants associated with height were included in our analysis as a fourth negative control to rule out a potential bias that could be introduced to our analysis by differences in the sizes of curated gene-sets. Our results indicate no statistically significant evidence for overrepresentation of AD or IBD-risk genes among the top neighbors of interrogated 3q29 interval genes. Only the top neighbors of *SENP5* (gene-set size = 713) showed a significant overlap with height-associated genes (overlap size = 30, adjusted *P* = 2.36E-02). Additionally, the top neighbors of *NRROS* (gene-set size = 68), which did not show an enrichment for known IDD, ASD, or SZ risk genes, exhibited a small but significant overlap with known PD-risk genes (overlap size = 3, adjusted *P* = 2.00E-02) (Fig. 3b).

Overall, 2 out of 64 hypergeometric tests indicated a significant overlap between the top neighbors of interrogated 3q29 interval genes and negative control gene-sets. In contrast, 19 out of 96 hypergeometric tests revealed a significant overrepresentation of SZ, ASD, and/or IDD-risk gene-sets among the same top neighbors, amounting to a proportion that is an order of magnitude larger than that of the negative controls (Fig. 3b). The substantial margin observed between these two enrichment ratios supports the high specificity and validity of our network-derived inferences for uncovering biology relevant to 3q29Del.

In summary, we identified 5,715 top neighbors of 3q29 interval genes, when combined across seven 3q29 modules. These top neighbors are predicted to function as direct interacting partners of 3q29 interval genes, participating in the same or overlapping biological pathways within the modular organization of molecular systems subserving the healthy functioning of the adult human PFC. Intriguingly, several 3q29 interval genes themselves were identified as top neighbors of other 3q29 interval genes, further suggesting functional convergence of subsets of genes within the 3q29 locus. Finally, our results revealed that *BDH1, CEP19*, *DLG1*, *FBXO45, PIGZ, RNF168, SENP5, UBXN7* and *WDR53* form strong co-expression-based ties with network partners that show a significant overlap with known SZ, ASD and/or IDD-risk genes curated from evidence-based literature (Fig. 3b). By leveraging the guilt by association principle, we prioritize these nine 3q29 interval genes, along with their 284 SZ, ASD, and/or IDD-related top neighbors from significant overlap tests as primary drivers of the major neuropsychiatric consequences of 3q29Del (Fig. 3b, Fig. 4a).

See Supp. Tables S2.1-3 for full lists of curated gene-sets, top neighbors, prioritized driver genes and detailed results of the overlap analysis.

**Disease-relevant driver genes prioritized by network analysis load onto key biological pathways linked to neuropsychiatric disorders.**

To formulate testable hypotheses about the key biological mechanisms linking the 3q29 locus to major neuropsychiatric phenotypes associated with 3q29Del syndrome, we interrogated whether the prioritized driver genes identified in our network analysis assemble into known biological pathways and processes that are annotated in major gene ontology databases. Functional enrichment analysis on the union of our 293 prioritized driver genes (Fig. 4a) revealed their significant overrepresentation in several key biological pathways and processes, some of which are specific to nervous system function, while others are core cellular processes that are non-specific to an organ system (Fig. 4b, 4c).

Specifically, our findings indicate enrichment of our prioritized driver genes in eight biological pathways annotated by the Reactome and KEGG databases. These are axon guidance (REAC:R-HSA-422475, adjusted *P* = 3.64E-03), post-translational protein modifications (REAC:R-HSA-597592, adjusted *P* = 5.24E-03), long-term potentiation (KEGG:04720, adjusted *P* = 7.29E-03), diseases of signal transduction (REAC:R-HSA-5663202, adjusted *P* = 1.00E-02), regulation of actin cytoskeleton (KEGG:04810, adjusted *P* = 1.17E-02), deubiquitination (REAC:R-HSA-5688426, adjusted *P* = 2.42E-02), chromatin organization (REAC:R-HSA-4839726, adjusted *P* = 3.32E-02), and diseases associated with glycosylation precursor biosynthesis (REAC:R-HSA-5609975, adjusted *P* = 4.06E-02). This analysis also revealed the enrichment of our prioritized driver genes for several fundamental biological processes annotated by the Gene Ontology Project (GO:BP), including chromosome organization (GO:0051276, adjusted *P* = 3.81E-09), histone modification (GO:0016570, adjusted *P* = 3.31E-08), cellular component morphogenesis (GO:0032989, adjusted *P* = 4.57E-06), regulation of organelle organization (GO:0033043, adjusted *P* = 6.40E-06), DNA metabolic process (GO:0006259, adjusted *P* = 3.83E-05), regulation of telomere maintenance (GO:0032204, adjusted *P* = 6.62E-05), neuron differentiation (GO:0030182, adjusted *P* = 1.88E-04), protein modification by small protein conjugation or removal (GO:0070647, adjusted *P* = 2.34E-04), neuron projection morphogenesis (GO:0048812, adjusted *P* = 4.46E-04), neurogenesis (GO:0022008, adjusted *P* = 1.89E-03), chemical synaptic transmission, postsynaptic (GO:0099565, adjusted *P* = 2.37E-03), post-embryonic development (GO:0009791, adjusted *P* = 2.66E-03), synapse organization (GO:0050808, adjusted *P* = 3.20E-03), protein acetylation (GO:0006473, adjusted *P* = 5.48E-03), cell surface receptor signaling pathway involved in cell-cell signaling (GO:1905114, adjusted *P* = 6.41E-03), and excitatory postsynaptic potential (GO:0060079, adjusted *P* = 8.97E-03). We hypothesize that the disruption of one or more of these biological pathways and processes, some of which have been demonstrated to be altered in idiopathic SZ and ASD [79, 80], lie on the casual pathway to neuropsychopathology in 3q29Del syndrome.

The top 20 biological processes and pathways enriched among our prioritized driver genes is shown in Fig. 4b. For clear illustration of our findings, we organized all identified GO:BP terms into a network of related functional annotation categories in Fig. 4c. Detailed results of this functional enrichment analysis, including a full list of prioritized drivers overlapping each identified gene ontology term are provided in Supp. Table S2.5.

In conclusion, prediction of novel gene-function and gene-disease associations is an important goal in computational biology, particularly for un- or under-studied territories of the human genome, such as the recurrent 3q29Del locus. These genes have been neglected, in part, due to attention bias in biomedical research that disproportionately concentrates on isolated interrogation of well-studied genes [84]. The network-based guilt-by-association approach used in this study is a promising strategy to rectify this skew and to advance our understanding of the full complement of the human genome and the full scope of genetic risk for severe mental illnesses in a systems biology framework.

### Availability of data & materials

We provide two multi-sheet xlsx files (Supp. Tables 1 and 2), containing further detailed results of our analysis. Corresponding R code is available upon request.

The GTEx data (release version 6) used for the analyses described in this manuscript were downloaded from the GTEx portal on 01/08/2019 (http://www.gtexportal.org/home/datasets/, file name: “GTEx_Analysis_v6_RNA-seq_RNA-SeQCv1.1.8_gene_rpkm.gct.gz”); dbGaP accession number: phs000424.v6.p1. The Genotype-Tissue Expression (GTEx) Project was supported by the Common Fund of the Office of the Director of the National Institutes of Health, and by NCI, NHGRI, NHLBI, NIDA, NIMH, and NINDS.

The BrainSpan data (Developmental Transcriptome Dataset) used to construct our test network were downloaded from the Allen Brain Atlas portal on 01/12/2019 (https://www.brainspan.org/static/download/, file name: “RNA-Seq Gencode v10 summarized to genes”); dbGaP accession number: phs000755.v2.p1.
