## Supplemental Figures for "Convergent and distributed effects of the schizophrenia-associated 3q29 deletion on the human neural transcriptome"

Sefik & Purcell et al.

Table of contents

Fig. S1. Tissue sample attributes and donor phenotypes of the GTEx dataset used for reference network construction…....................................................................................2
Fig. S2. Pre-processing of the reference dataset: outlier removal…........................................3
Fig. S3. Determination of the soft-thresholding power ($\beta$) in WGCNA……………………..……..4
Fig. S4. Determination of network reproducibility and module preservation in an independent test dataset……………………………………………………………..……………………....5
Fig. S5. Individual module-preservation and quality statistics underlying composite Zsummary scores……………………………………………………………………………………… 6-7
Fig. S6. Publication numbers for 3q29 genes and historic schizophrenia spectrum disorder candidate genes………………………………………………………………………..………8

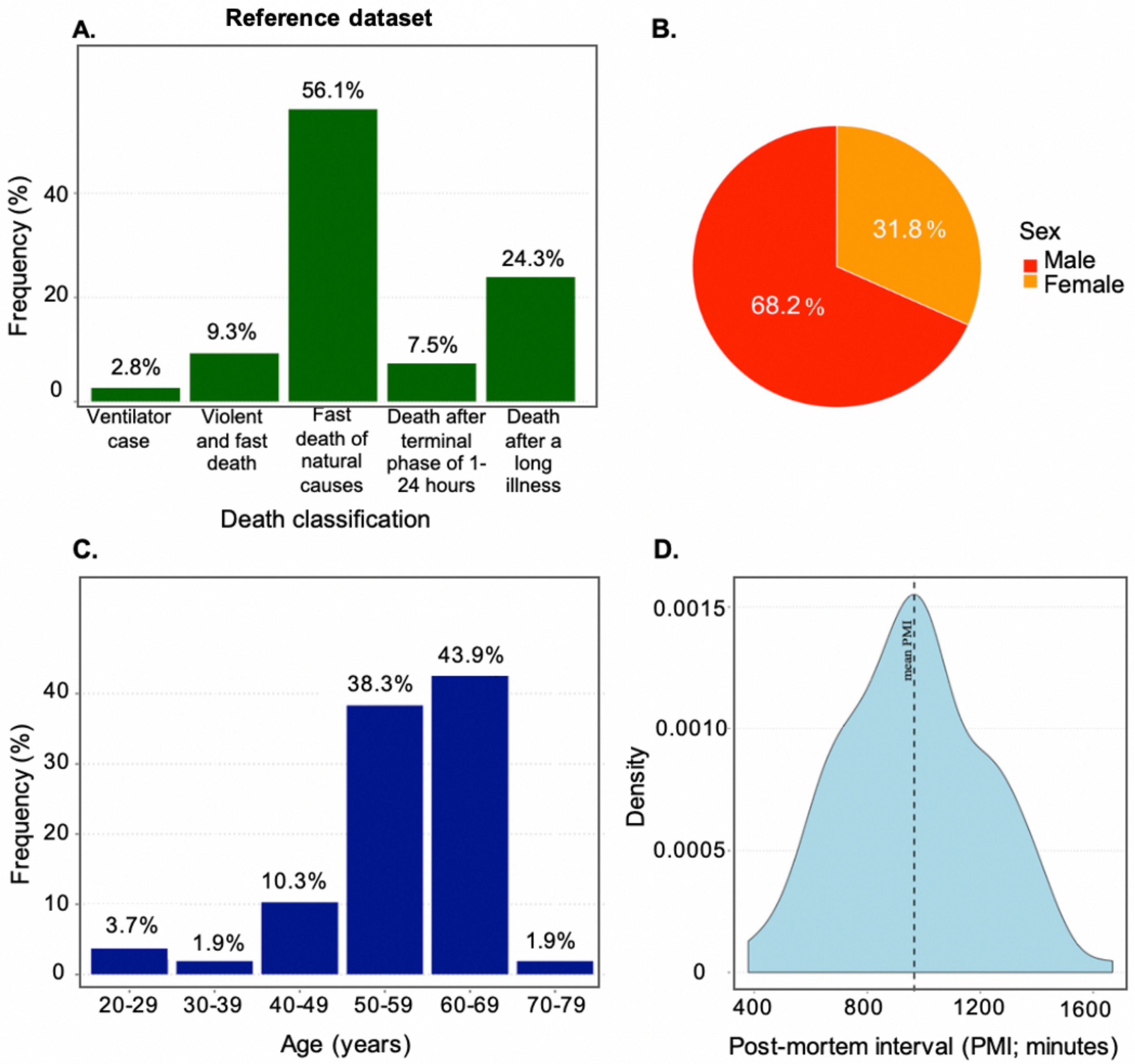


**Fig. S1. Tissue sample attributes and donor phenotypes of the GTEx dataset used for reference network construction (*N* = 107). (A)** Frequency distribution of donors' death-classification based on the 4-point Hardy scale. Majority of tissue samples (56.1%) were obtained from donors whose deaths were classified as "fast death of natural causes", which encompasses sudden (unexpected) deaths of people who had been reasonably healthy, following a terminal phase of <1 hour (e.g., sudden death from myocardial infarction). **(B)** Frequency distribution of donors' sex. Male:female ratio = 2.1:1.0. **(C)** Frequency distribution of donors' age-group based on 4-year intervals determined by the GTEx project. Majority of donors (82.2%) were in the age-range of 50-69 years at the time of death. **(D)** Sample density plot of post-mortem interval (PMI), indicating time elapsed between donor's death and final tissue stabilization. Mean sample PMI (sd) = 966.2 (254.3) minutes.

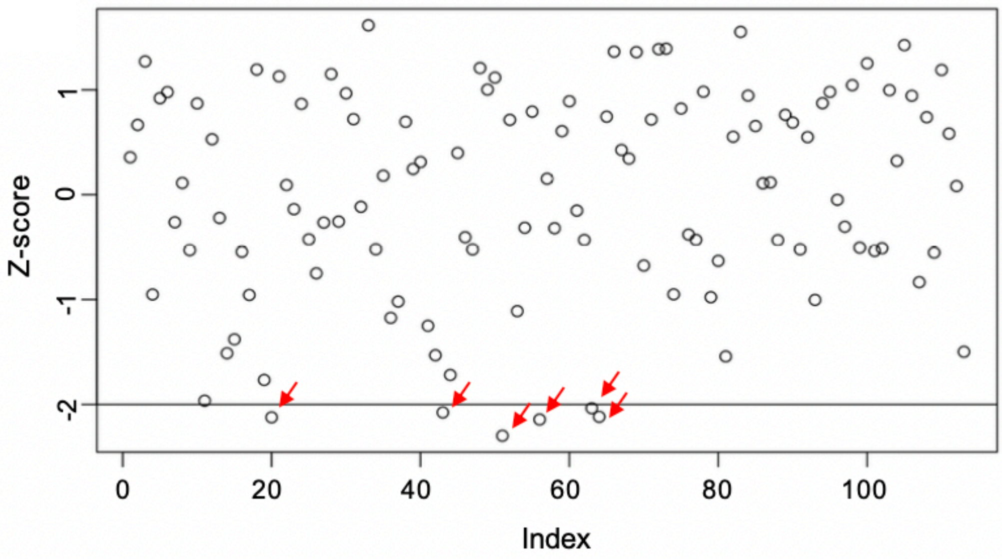


**Fig. S2. Pre-processing of the reference dataset: outlier removal.** 6 outlier samples (red arrows) were removed from the reference dataset (GTEx Project) to prevent an outlier-driven bias in gene co-expression network construction. Inter-sample correlation (ISC) was used as the statistical diagnostic for identifying samples with divergent gene expression profiles. ISC was defined as the Pearson's correlation between pairs of samples across the expression levels of all detected genes. Samples with a mean ISC greater than 2 standard deviations away (black line) from the mean of the sample-set were removed, bringing the sample size used for weighted gene co-expression analysis to 107.

A.

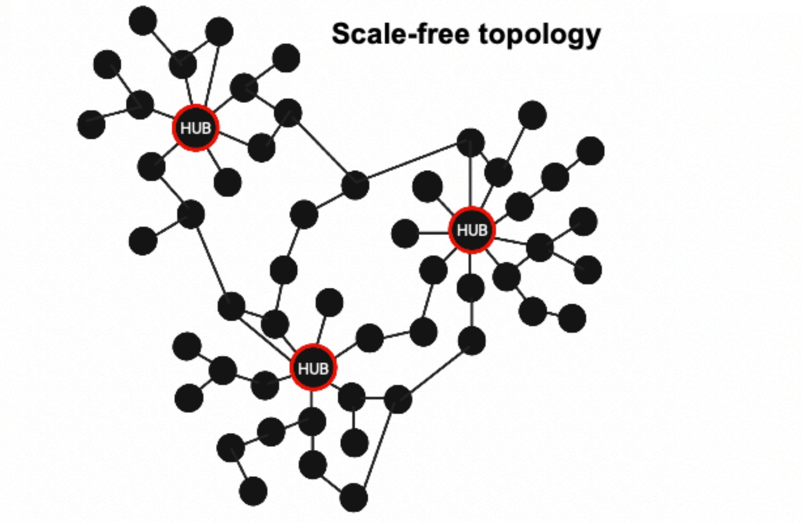


B.

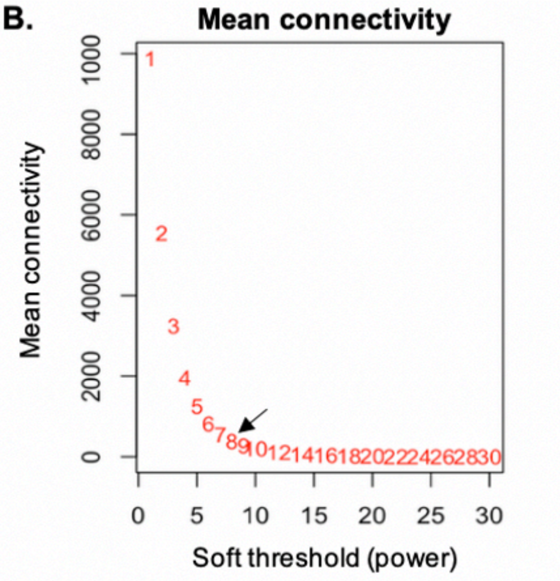


C.

| Power | SFT $R^2$ | Slope | Truncated $R^2$ | Mean k | Median k | Max k |
|---|---|---|---|---|---|---|
| 1 | 0.6 | 18.9 | 0.8 | 9890 | 9880 | 11100 |
| 2 | 0.5 | 7.0 | 0.9 | 5520 | 5460 | 7100 |
| 3 | 0.3 | 3.2 | 0.9 | 3200 | 3100 | 4740 |
| 4 | 0.0 | 0.5 | 0.8 | 1930 | 1810 | 3370 |
| 5 | 0.2 | -1.0 | 0.8 | 1200 | 1090 | 2520 |
| 6 | 0.6 | -1.6 | 0.9 | 773 | 674 | 1940 |
| 7 | 0.7 | -1.8 | 1.0 | 513 | 430 | 1540 |
| 8 | 0.8 | -1.8 | 1.0 | 351 | 281 | 1250 |
| 9 | 0.8 | -1.8 | 1.0 | 247 | 186 | 1020 |
| 10 | 0.9 | -1.8 | 1.0 | 178 | 125 | 854 |
| 12 | 0.9 | -1.7 | 1.0 | 98 | 60 | 613 |
| 14 | 0.9 | -1.8 | 1.0 | 58 | 30 | 459 |
| 16 | 0.9 | -1.8 | 1.0 | 36 | 16 | 353 |
| 18 | 0.9 | -1.7 | 1.0 | 24 | 9 | 278 |
| 20 | 0.9 | -1.7 | 1.0 | 16 | 5 | 222 |
| 22 | 0.9 | -1.7 | 1.0 | 11 | 3 | 180 |
| 24 | 0.9 | -1.7 | 1.0 | 8 | 2 | 148 |
| 26 | 0.9 | -1.7 | 1.0 | 6 | 1 | 123 |
| 28 | 0.9 | -1.7 | 1.0 | 4 | 1 | 103 |
| 30 | 0.9 | -1.7 | 1.0 | 3 | 0 | 87 |

D.

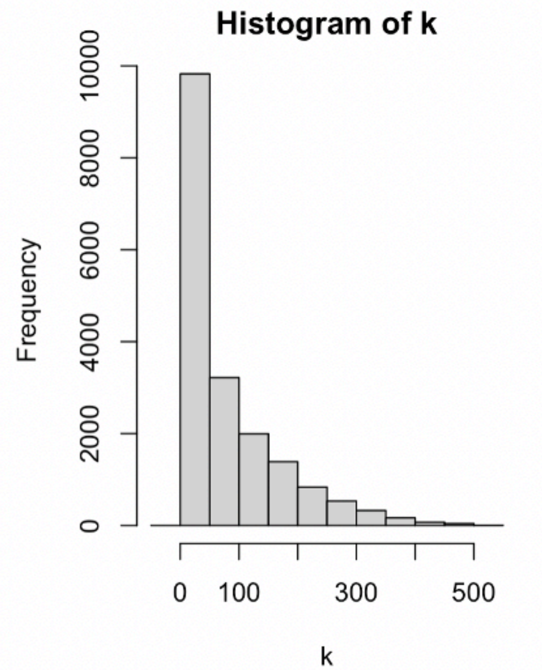


**Fig. S3. Determination of the soft-thresholding power ($\beta$) in WGCNA. (A)** Schematic of a representative scale-free network, whose degree distribution follows a power-law**:** most nodes have one or two connections but a few highly connected nodes (hub genes; marked by a red frame) have a large number of connections. Scale free topology (SFT) is a unifying property of biological networks in nature (Barabasi et al., 2004*). **(B)** Mean connectivity (k) as a function of different $\beta$ values. Mean k decreases as $\beta$ increases. The arrow marks the $\beta$ value used in this study. **(C)** List of SFT fitting indices for a wide range of $\beta$ values. The highlighted row indicates the $\beta$ used in this study. Given the necessary trade-off between SFT index $R^2$ and mean connectivity (k), a $\beta$ of 8 was identified as the lowest possible power yielding a degree distribution that results in approximate SFT ($R^2$ fit index = 0.8), while maintaining relatively high mean connectivity (mean k > 100), enabling the detection of modules and hub nodes. **(D)** Histogram of connectivity (k) distribution when a $\beta$ of 8 was chosen for defining the adjacency matrix. The frequency distribution of k shows a large number of lowly connected genes and a small number of highly connected genes, indicating that the resulting network follows the SFT criterion.

* Barabasi AL, Oltvai ZN. Network biology: understanding the cell's functional organization. *Nat Rev Genet* 2004; **5**(2)**:** 101-113.

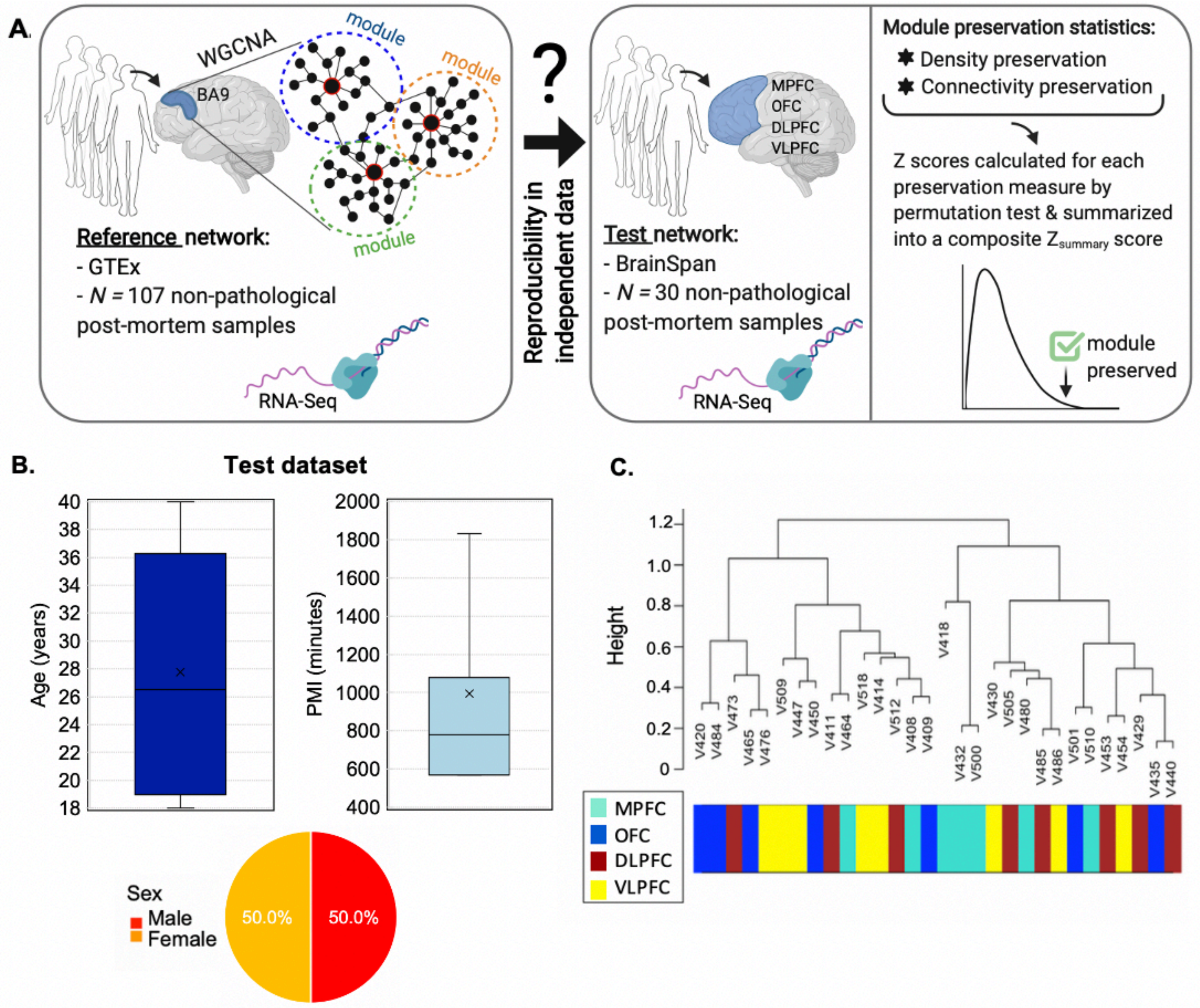


**Fig. S4. Determination of network reproducibility and module preservation in an independent test dataset. (A)** Illustration of the network reproducibility approach used in this study. Preservation of all network modules that were identified in the reference dataset (GTEx Project, N=107) was established in an independent, demographically comparable test dataset obtained from the BrainSpan Project (N=30). Both transcriptomic datasets were corrected for covariance mediated by sex, age and post-mortem interval (PMI). **(B)** Tissue sample attributes and donor phenotypes of the BrainSpan test dataset. The dark blue box-plot indicates the distribution of donors' age at death (in years). Median age = 27 years (range: 18-40 years). Mean age (sd) = 28 (8.4) years and corresponds to the cross mark. The light blue box-plot indicates the distribution of PMI in the test dataset. Median PMI = 780 minutes. Mean PMI (sd) = 994 (446.2) minutes. Pie chart indicates the percentage breakdown of donors' sex. Male:female ratio = 1:1. **(C)** Sample-level dendrogram of the test dataset plotted by hierarchical clustering of 30 non-pathological post-mortem samples obtained from four subregions of the prefrontal cortex (PFC) from male and female adults with no known history of neurological or psychiatric disorder. Clustering was conducted on normalized and residualized gene expression values for 18,339 protein-coding genes. Color bar below the dendrogram indicates tissue type corresponding to four subregions of the PFC that were pooled to derive the test dataset. The resulting dendrogram reveals no distribution bias associated with tissue-subtype in sample-level clustering patterns. OFC: orbital frontal cortex, DLPFC: dorsolateral PFC, VLPFC: ventrolateral PFC, MPFC: medial PFC.

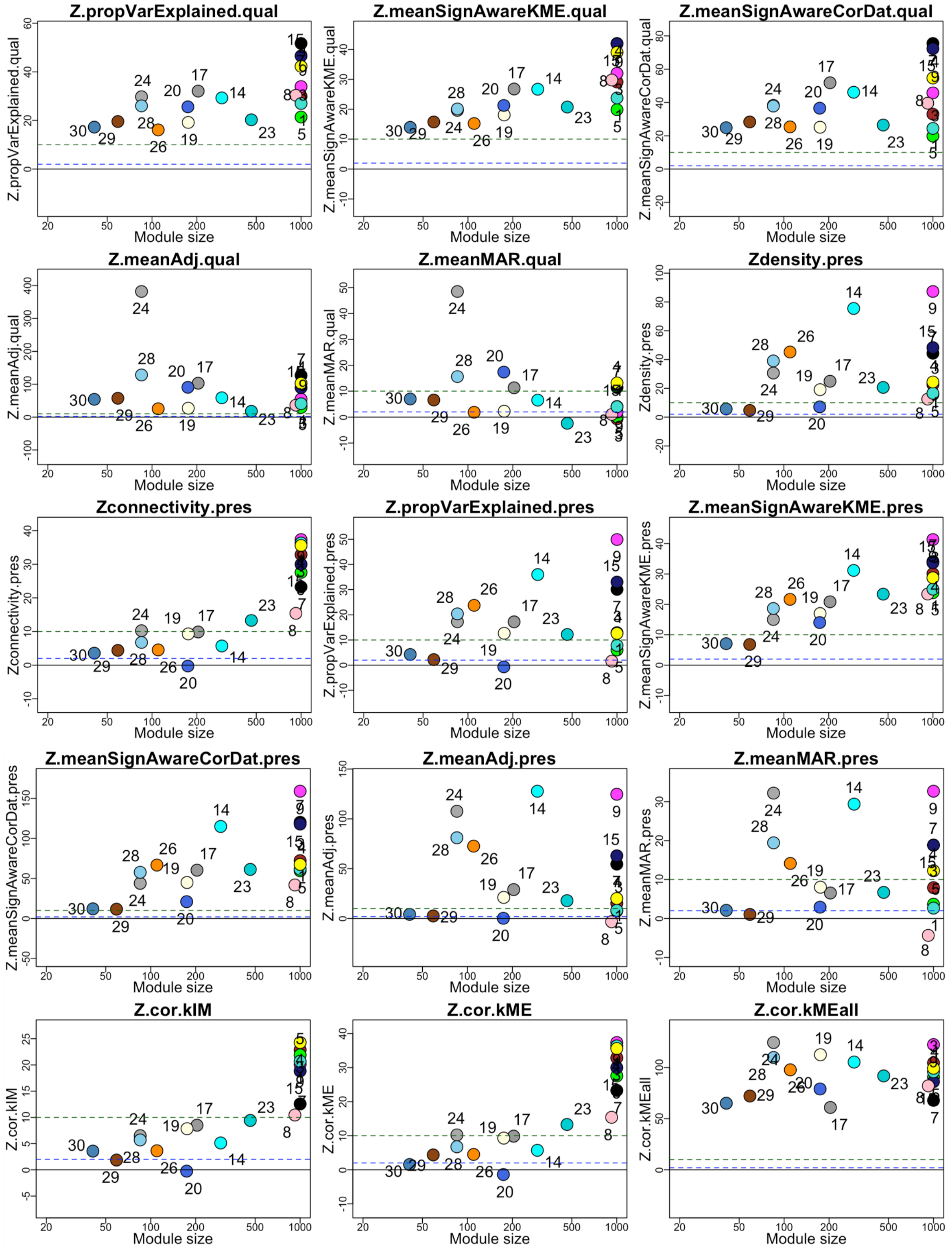


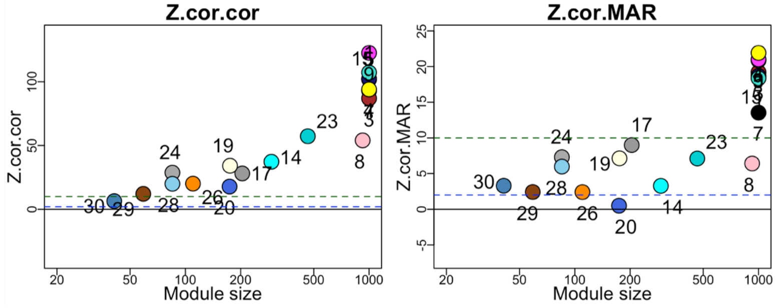


| module / color | label |
|---|---|
| turquoise | 1 |
| brown | 3 |
| dark turquoise | 23 |
| green | 5 |
| black | 7 |
| magenta | 9 |
| midnight blue | 15 |
| cyan | 14 |
| pink | 8 |

| | |
|---|---|
| grey60 | 17 |
| light yellow | 19 |
| royal blue | 20 |
| dark grey | 24 |
| dark orange | 26 |
| sky blue | 28 |
| saddle brown | 29 |
| steel blue | 30 |
| yellow | 4 |

**Fig. S5. Individual module-preservation and module-quality statistics underlying composite Zsummary scores.** Multiple density-based and connectivity-based module-preservation statistics were assessed to determine the distinct properties of network structure preserved between the reference and test networks. In line with the composite $Z_{summary}$ statistic, individual module-preservation statistics mostly converged on the finding that 1) nodes sharing the same module in the reference network remain highly connected in the test network and 2) connectivity patterns between nodes underlying the reference network remain similar in the test network. Similarly, multiple module-quality based statistics were assessed to determine how distinct individual modules were from all other modules in the reference network. In line with the composite $Z_{summary}$ statistic, the majority of the evaluated module-quality statistics indicate robust module definitions across networks created from random splits of the original reference data. Permutation tests were performed to adjust the observed preservation and quality statistics of each module for random chance by defining Z statistics. Zstatistic < 2 (blue dotted line): no evidence for preservation/quality; 2 < Zstatistic < 10 (green dotted line): moderate evidence for preservation/quality; Zstatistic > 10: strong evidence for preservation/quality. Refer to Langfelder et al. (2017)* for a detailed description of the individual module preservation and quality statistics plotted above. Both numeric and color-based labels were used to mark individual modules. The coding system corresponding to module labels is provided in the bottom right corner of the figure; labels highlighted in yellow indicate the 7 modules harboring 3q29 interval genes.

* Langfelder P, Luo R, Oldham MC, Horvath S. Is My Network Module Preserved and Reproducible? *Plos Computational Biology* 2011; **7**(1).

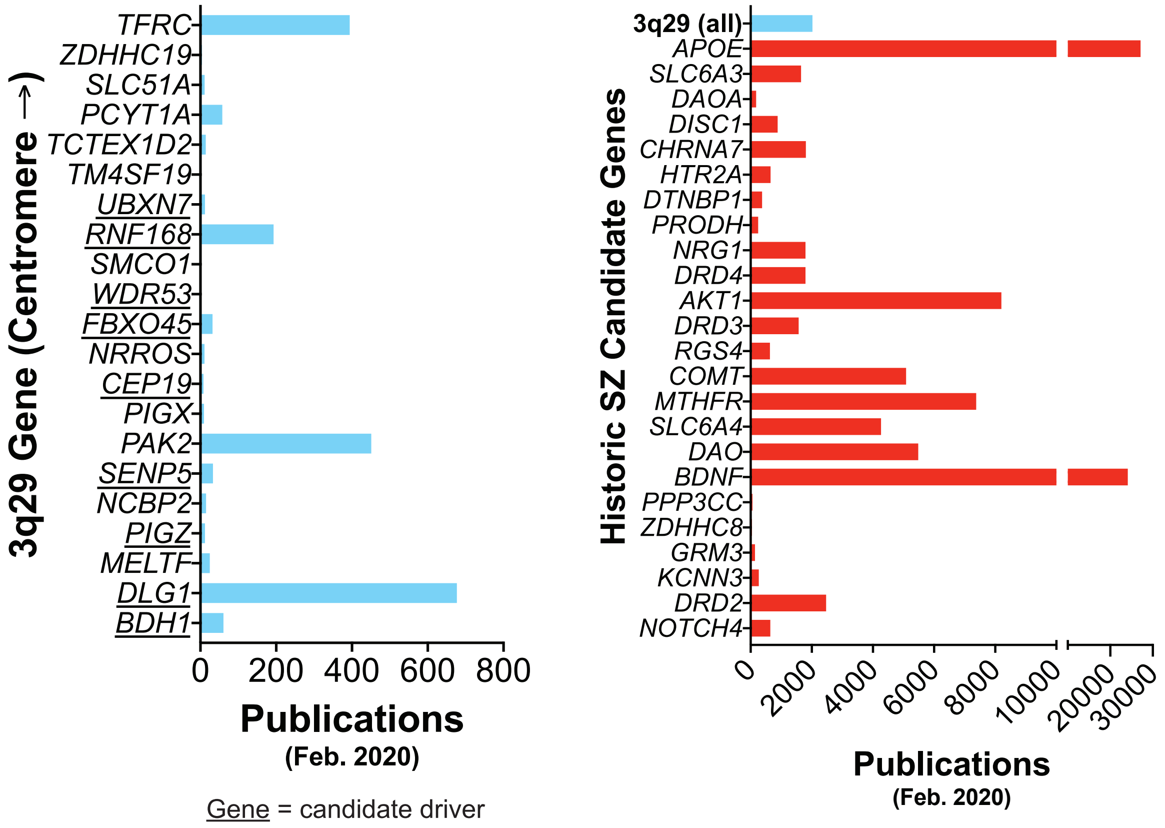


**Fig. S6. Publication numbers for 3q29 genes and historic schizophrenia spectrum disorder candidate genes.** The total number of PubMed articles retrieved with a search for the symbol of each gene is shown above. Notably, of the prioritized driver genes (underlined) identified in this study (left), only *DLG1* has a publication history that compares favorably with historic schizophrenia spectrum disorder (SZ) candidate genes (right) that were typically identified by genetic linkage and association studies. The majority of 3q29 interval genes remain understudied.
